# μ Opioid Receptors Modulate Action Potential Kinetics and Firing Frequency in Neocortical Interneurons

**DOI:** 10.1101/2020.11.20.391508

**Authors:** Adrian P Dutkiewicz, Anthony D. Morielli

## Abstract

The endogenous opioid system of the cerebral cortex is an important feature of antinociception and reward valuation through its modulation of inhibitory neocortical interneurons. Dysregulation of this system, through disease or drugs, disrupts the reward system and contributes to eating and mood disorders, impulsive actions, and addiction. Impulsive behaviors can be induced experimentally through infusion of the μ opioid receptor specific agonist [D-Ala2, N-Me-Phe4, Gly5-ol]-Enkephalin (DAMGO) into the frontal cortex in animal models. The mechanism involves increased potassium channel function, which suppresses neocortical interneuron activity. However, much of the data on the effect of this receptor on ion channels have been derived from noncortical μORs, and the identity and effects of the ion channels that the μOR targets in neocortical neurons have not been thoroughly investigated. Based on previous experiments by other labs, we hypothesized that the μOR could activate α-dendrotoxin (αDTX) sensitive channels (Kv1.1, Kv1.2, and Kv1.6 subunits) to exert its inhibitory effects in cortical interneurons. This, in turn, is expected to confer a variety of effects on passive and active electrical properties of the cell. We performed patch-clamp electrophysiology to examine the electrophysiological effects of μORs in cultured neocortical interneurons. We found that a range of features among the 54 membrane and action potential properties we analyzed were modulated by μORs, including action potential kinetics and frequency. The Kv1.1, Kv1.2, and Kv1.6 inhibitor αDTX reversed some effects on action potential frequency, but not effects on their kinetics. Therefore, μORs in neocortical interneurons influence αDTX-sensitive channels, as well as other channels, to modulate action potential kinetics and firing properties.

## INTRODUCTION

The endogenous opioid system of the cerebral cortex mediates antinociception and reward valuation (Choi et al., 2016; Ong, Stohler, & Herr, 2019; Qiu, Wu, Xu, & Sackett, 2009; Zubieta et al., 2001). Dysregulation of this system is believed to contribute to pathological and compulsive behaviors such as eating disorders, pathological gambling, and drug-seeking (Ashok, Myers, Reis Marques, Rabiner, & Howes, 2019; Bencherif et al., 2005; Giuliano & Cottone, 2015; Joutsa et al., 2018; Mick et al., 2016; Mitchell et al., 2012; Qu, Huo, Huang, & Tang, 2015; Zubieta et al., 1996). Experimentally, impulsive behaviors and binge-eating can be induced through infusion of the μ opioid receptor (μOR) specific agonist [D-Ala2, N-Me-Phe4, Gly5-ol]-Enkephalin (DAMGO) into the frontal cortex of animal models (Mena, Sadeghian, & Baldo, 2011; Selleck et al., 2015). Blockade of μORs with naltrexone inhibits compulsive behaviors (Bartus et al., 2003; Blasio, Steardo, Sabino, & Cottone, 2014). These aberrancies are believed to result from the disruption of activity within cortical networks by μORs (Haider, Duque, Hasenstaub, & McCormick, 2006; Whittington, Traub, Faulkner, Jefferys, & Chettiar, 1998; Zhang et al., 2019). More specifically, μORs appear to dysregulate cortical networks by suppressing GABAergic activity and thereby altering the balance of excitation and inhibition in the cortex (Lewis, Curley, Glausier, & Volk, 2012; Shi et al., 2020; Volk, Radchenkova, Walker, Sengupta, & Lewis, 2012). The μOR is believed to suppress GABAergic signaling through its expression primarily on cortical interneurons. This leads to overactivity of the targets of their inhibition, the glutamatergic Pyramidal Neurons (PNs) (Drake & Milner, 1999, 2002, 2006; Férézou et al., 2007; Huo et al., 2005; Madison & Nicoll, 1988; Stumm, Zhou, Schulz, & Höllt, 2004; Taki, Kaneko, & Mizuno, 2000; Witkowski & Szulczyk, 2006; Zieglgansberger, French, Siggins, & Bloom, 1979). However, some research suggests that μOR may activate PNs directly as well (Rola, Jarkiewicz, & Szulczyk, 2008; Schmidt et al., 2003; Witkowski & Szulczyk, 2006). Immunohistochemical experiments have found high rates of expression of μOR in interneurons that express vasoactive-intestinal peptide (VIP+) (Taki et al., 2000). Electrophysiological and sc-PCR data in neocortical neurons have supported this finding (Férézou et al., 2007) and have also implicated neurogliaform neurons as expressing this receptor (Férézou et al., 2007; Lafourcade & Alger, 2008; McQuiston, 2008). Evidence for μOR expression in somata of parvalbumin-positive (PV+) interneurons of the hippocampal formation is fairly clear (Bartos & Elgueta, 2012; Bausch et al., 1995; Drake & Milner, 1999, 2002, 2006; Stumm et al., 2004; Torres-Reveron et al., 2009), however perisomatic expression of this receptor in neocortical PV+ neurons has been investigated but, to our knowledge, has not been reported (Férézou et al., 2007; Taki et al., 2000), though recent evidence shows that μORs are sometimes found in PV+ axon terminals of the frontal cortex (Jiang et al., 2019; Lau, Ambrose, Thomas, Qiao, & Borgland, 2020) and insular cortex (Yokota et al., 2016). Therefore, neocortical VIPergic, neurogliaform, and some PV+ interneurons are generally believed to express μORs – though not necessarily in their somata.

In addition to the uncertainty surrounding the μOR’s localization, there is also some mystery surrounding its mechanism by which μORs affect interneuron activity. The μOR is a G-protein coupled receptor and has been shown to activate potassium-conducting inwardly-rectifying K (GIRK) channels (Henry, Grandy, Lester, Davidson, & Chavkin, 1995; Ikeda, Kobayashi, Kumanishi, Niki, & Yano, 2000; Loose & Kelly, 1990; Marker, Lujan, Loh, & Wickman, 2005). Regulation of GIRK channels and hyperpolarization are characteristic features of various ORs (Ikeda, Yoshii, Sora, & Kobayashi, 2003). Agonist-induced hyperpolarization has been found in cortical μOR+ neurons (Férézou et al., 2007; Glickfeld, Atallah, & Scanziani, 2008; Madison & Nicoll, 1988; Tanaka & North, 1994) non-cortical μOR+ neurons (Grudt & Williams, 1993; Harris & Williams, 1991; Johnson & North, 1992; Kelly, Loose, & Ronnekleiv, 1990; Lagrange, Ronnekleiv, & Kelly, 1994; Lagrange, Rønnekleiv, & Kelly, 1995; Loose & Kelly, 1990; North, Williams, Surprenant, & Christie, 1987; Sugita & North, 1993), and is also a feature of other opioid receptors as well (Chieng & Christie, 1994; Chiou & Huang, 1999; Grudt & Williams, 1993; Johnson & North, 1992; Loose & Kelly, 1990; Loose, Ronnekleiv, & Kelly, 1990; Madison & Nicoll, 1988; North et al., 1987; Sugita & North, 1993). However, hyperpolarization does not always occur in response to DAMGO (Faber & Sah, 2004; Wimpey & Chavkin, 1991), possibly due to an incomplete overlap between GIRK channel expression and μORs, which may be the case in the cortex (Bausch et al., 1995). Additionally, reductions in spontaneous APs with μOR-agonism is also found in various parts of the brain, though not all μOR+ neurons fire spontaneously (Loose & Kelly, 1990; Mitrovic & Celeste Napier, 1998; Ponterio et al., 2013). Hyperpolarization and decreased spontaneous activity are both commonly found in response to the activation of μOR and other opioid receptors, and the two effects often coincide (Chiou & Huang, 1999; Elghaba & Bracci, 2017; Kelly et al., 1990; Loose & Kelly, 1990; North et al., 1987). While it is possible that hyperpolarization could induce a decrease in spontaneous APs by increasing the distance to V_m_ threshold for action potentials, some studies in cortical neurons have found reductions in spontaneous APs with only small accompanying hyperpolarization (Krook-Magnuson, Luu, Lee, Varga, & Soltesz, 2011; M. E. Sheffield et al., 2013; M. E. J. Sheffield, Best, Mensh, Kath, & Spruston, 2011; Suzuki, Tang, & Bekkers, 2014). Therefore, the μOR may induce hyperpolarization of varying magnitude along with reductions in tonic APs, and one effect could occur independently of the other.

In addition to hyperpolarization and reductions in spontaneous APs, research from other parts of the brain suggests that μORs modulate αDendrotoxin-sensitive channels; αDTX inhibits Kv1.1, Kv1.2, and Kv1.6 channels (Ponterio et al., 2013). Experiments have found that μORs modulates αDTX-sensitive channels in the basolateral amygdala (Finnegan, Chen, & Pan, 2006), periaqueductal gray (Vaughan, Ingram, Connor, & Christie, 1997), as well as thalamocortical terminals within the frontal cortex (Lambe & Aghajanian, 2001). Although these channels are known to be expressed by several families of neocortical interneurons (Casale, Foust, Bal, & McCormick, 2015; Goldberg et al., 2008; Golding, Jung, Mickus, & Spruston, 1999; Li et al., 2014; Porter et al., 1998), they have not been directly investigated for mediating the μOR’s inhibitory effect in the neocortical interneurons. We therefore predicted that μORs activate αDTX-sensitive channels to inhibit neocortical interneurons.

Several studies have investigated the role of the αDTX-sensitive channels by analyzing its effects on action potentials. While Kv1.1, Kv1.2, and Kv1.6 channels have been shown to modulate spike frequency and AP threshold, (Bekkers & Delaney, 2001; Faber & Sah, 2004; Finnegan et al., 2006), and some studies have suggested that they contribute to AP shape as well. Specifically, researchers have found that αDTX-sensitive channels hasten repolarizations and shorten durations of APs (Geiger & Jonas, 2000; Pathak, Guan, & Foehring, 2016), including in neocortical interneurons (Casale et al., 2015). We therefore predicted that part of the mechanisms by which the μOR act in neocortical interneurons is through one or a combination of these αDTX channel-mediated electrophysiological effects. To investigate this, we cultured neocortical neurons and performed patch-clamp electrophysiology on interneurons in current-clamp mode to stimulate and measure their APs. To measure the kinetics of APs, we created Python scripts to measure 54 membrane properties, AP kinetic properties, and ratios. We were primarily interested in 7 properties that have previously been implicated in mediating DAMGO or αDTX effects in neurons. We predicted that DAMGO and αDTX would modulate (in opposite polarity) resting membrane potential, AP threshold, number of evoked APs, interspike interval, AP halfwidth, maximum repolarization rates, amplitude of afterhyperpolarizations.

## RESULTS

### Identification of DAMGO-responding neurons

Cortical μOR is expressed only in certain subcategories of interneurons; some studies have estimated that only around 3-5% of all neocortical neurons, or 15-25% of GABAergic interneurons, express μORs (Férézou et al., 2007; M. C. Lee, Cahill, Vincent, & Beaudet, 2002; Taki et al., 2000), though some have reported as many as 2/3rds of prefrontal GABAergic neurons respond to μOR-agonism (Witkowski & Szulczyk, 2006). To identify and target GABAergic interneurons in culture, we transformed neurons with an AAV (see Methods) that drove expression of the red fluorescent protein mRuby2 under the interneuron-specific Dlx5/6 enhancer (Batista-Brito, Machold, Klein, & Fishell, 2008; de Lombares et al., 2019; Fazzari et al., 2010).

Neocortical interneurons generally only constitute between 10-25% of all neocortical neurons (with the balance being glutamatergic Pyramidal Neurons), depending on the model and methods used to measure the proportions (Beaulieu, 1993; Jones, 1993; Meyer et al., 2011; Ren, Aika, Heizmann, & Kosaka, 1992). To determine whether proportions of interneurons occurring in the culture model used here fell within this range, we manually counted the number of red and nonred neurons across 110 images in 3 independent cultures. We found that red neurons constituted 432 out of 2428 neurons (17.8%). In a second approach, we utilized a Python script that automatically counted the neurons. This automated approach gave similar results, with red neurons constituting 1037 out of 6461 neurons (16.1%). With either counting method, our estimations of the proportions of interneurons to non-interneurons in these dissociated neuronal cultures are typical for previous reports of intact or sliced animal neocortex.

Overall, the DAMGO-exposed neurons (*N* = 55, *M* = 1.03, *SD* = 0.05) were significantly more (*t*(74) = 3.14, *p* = 0.001) hyperpolarized than saline controls (*N* = 21, *M* = 0.99, *SD* = 0.99), which tended to slightly depolarize (Figure 3c). However, the μOR is only expressed by a subset of cortical interneurons, and therefore was expected to be represented in only some of the red-fluorescent Dlx5/6-mRuby2 neurons that we recorded from. We anticipated that most neocortical interneurons would be unresponsive to DAMGO. Combining data from all interneurons would thus obscure the effects of DAMGO (3 μM) and αDTX (100 nM) in neurons that did express had μORs. We therefore screened interneurons for responsiveness to DAMGO prior to subsequent analyses. Hyperpolarization is a common response to DAMGO in neurons expressing opioid receptors (Chieng & Christie, 1994; Chiou & Huang, 1999; Grudt & Williams, 1993; Johnson & North, 1992; Loose & Kelly, 1990; Loose et al., 1990; Madison & Nicoll, 1988; North et al., 1987; Sugita & North, 1993), and in particular those expressing μORs (Férézou et al., 2007; Glickfeld et al., 2008; Harris & Williams, 1991; Kelly et al., 1990; Lagrange et al., 1994; Lagrange et al., 1995; Madison & Nicoll, 1988; Tanaka & North, 1994). We therefore first identified responders by detecting a hyperpolarizing change in resting membrane potential (RMP) after exposure to DAMGO (Figure 3). By comparing the DAMGO-exposed group’s changes in RMP to the changes in saline controls, we established a cutoff level to separate neurons most affected by DAMGO. We refer to the DAMGO-exposed neurons above this cutoff as “hyperpolarizing responders” (H-responders, *N* = 11). These H-responders hyperpolarized significantly (*t*(20) = 7.61, *p* < 0.001) when they (*N* =11, *M* = 1.10, *SD* = 0.03) were compared to a group of random saline controls (*n* = 11, 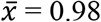, *s* = 0.04).

During the recording process, we observed that only 3 of the H-responders were firing spontaneously at rest and, not surprisingly, their rate of spontaneous APs decreased after DAMGO. However, we observed that a sub-population of interneurons in which DAMGO failed to produce an above cut-off level of hyperpolarization nevertheless underwent reductions in spontaneous APs after DAMGO. We found that 50.0% (22 out of 44) of neurons that fell below the H-responder cut-off level exhibited spontaneous AP activity before DAMGO (Figure 4). Application of DAMGO resulted in a dramatic reduction in the number of spontaneous APs in 45.5% (10 out of 22) of those neurons. We define this group of spontaneous AP DAMGO-responders (S-responders) by using an arbitrary cutoff of a 50% reduction in spontaneous APs after DAMGO (*N* = 10). Spontaneous APs were measured during a noncontiguous period (cumulative of 60s) in between current pulses.

Figure 4 depicts the fold change in number of spontaneous APs (slot 2/slot 1) in the saline group (*N* = 11) and the DAMGO-exposed group (*N* = 22). Because a saline control dropped spontaneous APs to zero, we could not use a “real datapoint” cutoff as we did with the H-responders. We therefore established a 50% reduction (0.50; at the vertical dotted line in Figure 4c) as a cutoff point for S-responders and nonresponders. All DAMGO-exposed neurons to the left of that cutoff were considered to be “S-responders” (*N* = 10). We did not analyze the effect of DAMGO on spontaneous APs beyond counting them, because 8 out of 10 S-responders had simply stopped firing spontaneous APs after DAMGO. Collectively, 21 out of 55 (38.2%) neurons fell into at least one of the two responder categories. Hyperpolarizing responders had an average RMP of −54.0 mV (range −50.1 to −62.9 mV). The H-responders had, on average, hyperpolarized by −5.6 mV (range −3.5 to −8.6 mV) during the 80s period from slot 1 (saline) to slot 2 (DAMGO). Most H-responders were silent at rest, with only three firing spontaneous APs. On average the S-responders fired around 2.8 APs/s before DAMGO when unstimulated. This ranged from 0.17 APs/s to 12.1 APs/s. After DAMGO, 8 out of 10 S-responders simply stopped firing spontaneous APs, however 2 continued to fire spontaneous APs at lower rates after DAMGO. A neuron that was firing spontaneous APs at 12.1 APs/s reduced its spiking to 5.75 APs/s after DAMGO, whereas the remaining the remaining neuron underwent a nearly four-fold reduction in spontaneous APs (Figure 4d).

To further explore whether the H and S-responder classes of DAMGO-responsive cells represent distinct cell populations, we determined the pre-drug spiking pattern based on the Petilla Interneuron Nomenclature Group’s suggested categorization scheme (The Petilla Interneuron Nomenclature, 2008). Categorization of H and S-responders was determined for based on their firing pattern during the 1s suprathreshold current application (Figure 5). Although we characterized their spiking pattern around their threshold V_m_ for the first AP, we observed that discharge patterns could change at higher current applications. For instance, fastspiking neurons typically had low firing rates around threshold, but increased at more-depolarizing current steps, which has been reported before (Golomb et al., 2007).

We found that S-responders and non-responders were roughly equivalent in their distribution of firing patterns. H-responders, in contrast, were notable in having a relatively high frequency of Adapting neurons, a relatively low frequency of non-adapting neurons and a lack of fast-spiking neurons. Collectively, these data suggest that the μOR effects might be heterogeneous and could vary between neuron populations.

### DAMGO effects on action potential activity and kinetics

Modulation of AP kinetics can affect neurotransmitter release by influencing the kinetics and amplitude of calcium influx (Yang & Wang, 2006). Features of AP kinetics are shaped by multiple types of ion channels (Rudy et al., 2009). We therefore evoked APs before and after DAMGO and αDTX to assess a potential role of μOR in regulating αDTX-sensitive ion channels as a means of governing AP kinetics.

We analyzed 54 AP parameters, ratios, and membrane properties (Tables 7 and 9). Of those, seven have been reported to be affected by activation of μORs or αDTX in other studies, though not necessarily both. Hyperpolarization was the first measure, which is often found with DAMGO stimulation (Loose et al., 1990). Secondly, αDTX-sensitive channels are shown to modulate V_m_ threshold for an AP (AP threshold) (Bekkers & Delaney, 2001; Glazebrook et al., 2002; Kirchheim, Tinnes, Haas, Stegen, & Wolfart, 2013; Pathak et al., 2016), though the μOR may not change it (Bekkers & Delaney, 2001; Faber & Sah, 2004; Glazebrook et al., 2002; Kirchheim et al., 2013; Pathak et al., 2016). Both the μOR and αDTX-sensitive channels also suppresses evoked AP discharges, and so we investigated whether it decreases the number of evoked APs and increases their temporal spacing (interspike interval; ISI) (Faber & Sah, 2004; Mo, Adamson, & Davis, 2002). The AP halfwidth (duration) and the AP maximum repolarization rate were also measured because there is some evidence to support their modulation by αDTX-sensitive channels (W. Wang, Kim, Lv, Tempel, & Yamoah, 2013). Finally, since many voltage-sensitive K channels contribute to the afterhyperpolarization (AHP), we measured the magnitude of afterhyperpolarizations as well (W. Wang et al., 2013).

### DAMGO alters AP waveform and pattern of evoked APs in H-responders

To reduce family-wise errors from running sets of 7 analyses simultaneously, we first tested the combined 7 dependent variables (RMP, AP threshold, interspike interval, number of evoked APs, AP halfwidth, maximum repolarization rate, and afterhyperpolarization amplitude) in the H-responders (*N* = 11) versus the saline controls (*N* = 21) for a significant Slot*Group interaction (see Table 1) in a mixed-model repeated measures MANOVA. When the Slot*Group interaction was significant, we tested the changes in the dependent variables individually.

**Table 1.** Control and Drug Groups. We included 2 groups in the experimental design; a saline-only control and the experimental group that received the sequence of drugs. In both cases, slot 1 recordings are always pre-drug conditions. Slot 2 is synonymously referred to as “post-DAMGO” or as “pre-αDTX” depending on whether we are testing for the effects of (3 μM) DAMGO or (100 nM) αDTX in that particular analysis; the αDTX was always delivered in a background of DAMGO. To detect drug effects, we analyzed the changes between 2 timeslots in the drug group and compared it to the changes in the saline group; slot 1 versus slot 2 for DAMGO effects, and slot 2 versus slot 3 for (100 nM) αDTX effects.

|  | Group | Within-Subjects Factor (Slot) |  |  |
| --- | --- | --- | --- | --- |
|  |  | Slot 1 | Slot 2 | Slot 3 |
| Between-Subjects Factor (Group) | Saline (N = 21) | Saline | Saline | Saline |
| | Drug Sequence (N = 55) | Saline | DAMGO | αDTX+DAMGO |

We found that the H-responders had statistically significant (*F*(7,20) = 19.32, *p* < 0.001, *Wilks’ λ* = 0.871) Slot*Group interaction in the 7 combined variables that we were tested, which suggested a possible DAMGO effect in those parameters. We therefore carried out specific analyses of RMP, AP threshold, number of evoked APs, interspike interval, AP halfwidth, max repolarization rate, and afterhyperpolarization magnitude in H-responders.

The αDTX-sensitive channels can modulate V_m_ threshold for an AP, however the μOR does not necessarily change this property (Faber & Sah, 2004; Finnegan et al., 2006). We analyzed their change in V_m_ threshold for the first AP (Figure 6a,b,c). We found no significant difference (*t*(20) = 0.42, *p* = 0.338) in the change in AP threshold for the H-responders (M = 0.99, *SD* = 0.07) versus the saline controls (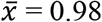, *s* = 0.07) after DAMGO. This shows that neocortical μORs do not modulate AP threshold in the H-responders.

We then investigated the patterns of AP discharges to determine whether how DAMGO had influenced the number of evoked APs and their temporal spacing between APs (Figure 6). These H-responders (*N* = 10, *M* = 1.35, *SD* = 0.36) had significantly increased interspike intervals (*t*(18) = 3.01, *p* = 0.004) when compared to the saline-only controls (*n* = 10, 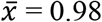, *s* = 0.12). The H-responders (*N* = 11, Mdn = 0.98) also had significantly reduced numbers of evoked action potentials (*U* = 12, *p* < 0.001) after application of DAMGO when compared to saline controls (*n* = 11, Mdn = 0.98). These data show that DAMGO reduced the number of APs evoked by suprathreshold currents.

We next analyzed the H-responder group to determine whether AP waveform was being altered by DAMGO (Figure 7). Among hyperpolarizing DAMGO-responders, we found several significant changes in action potential waveforms against the saline-only controls; we found significant reductions in AP halfwidth (*U* = 11, *p* < 0.001) in the DAMGO H-responders (Mdn = 0.91) versus the saline controls, which trended slightly towards widening APs over that time period (Mdn = 1.05). The 11 DAMGO H-responders had increased their maximum repolarization rates (*M* = 1.29, *SD* = 0.44) significantly (*t*(20) = 2.91, *p* = 0.004) compared to the saline controls (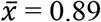, *s* = 0.11) which trended towards slower repolarizations by slot 2. These results show that DAMGO was hastening the maximum rates of AP repolarization, and shortening the duration of the APs. Lastly, we compared the magnitude of afterhyperpolarizations (AHPs) in the saline controls (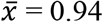, *s* = 0.07) versus the H-responders (*M* = 1.10, *SD* = 0.30) and found that DAMGO had significantly increased (*t*(20) = 1.78, *p* = 0.045) the AHPs in the H-responders.

We classified the H-responders by spiking pattern after DAMGO to determine if μORs could change the pattern of discharges, since there were changes in evoked AP number and ISI in these neurons (Table 4). Although most H-responders had a stable spiking pattern after DAMGO, we did notice a few changes. Three H-responders had shifted spiking categories, however the majority (8 out of the 11) H-responders had remained in the same spiking pattern after DAMGO. In 2 out of 3 cases where the H-responder shifted pattern, the neurons transitioned into an Adapting patten, which was consistently the prevailing spiking pattern in H-responders.

### αDTX counteracts μOR-agonism in H-responders

Having found that DAMGO was altering AP waveform and pattern of discharge, particularly in hyperpolarizing DAMGO responders, we investigated whether αDTX-sensitive channels were involved in this effect; μOR elsewhere in the brain has previously been found to modulate an α-Dendrotoxin-sensitive channels (Faber & Sah, 2004; Finnegan et al., 2006). We therefore exposed the neurons to a combination of DAMGO + αDTX (3 μM and 100 nM, respectively) after their prior exposure to (3 μM) DAMGO to determine if the addition of αDTX could reverse the effects. We once again compared this drug group against saline-only recordings to account for stochastic effects and the effects of repeated stimulation, and thereby detect the true impact of αDTX in a background of DAMGO. For the following comparisons we also isolated the change by dividing the post αDTX+DAMGO/pre αDTX-DAMGO; slot 3/slot 2. In this case, the post-DAMGO (slot 2) measurements that had been used as the “after-DAMGO” recording were used here as the “pre-αDTX+DAMGO” recording (see Figure 2).

**Figure 1.**
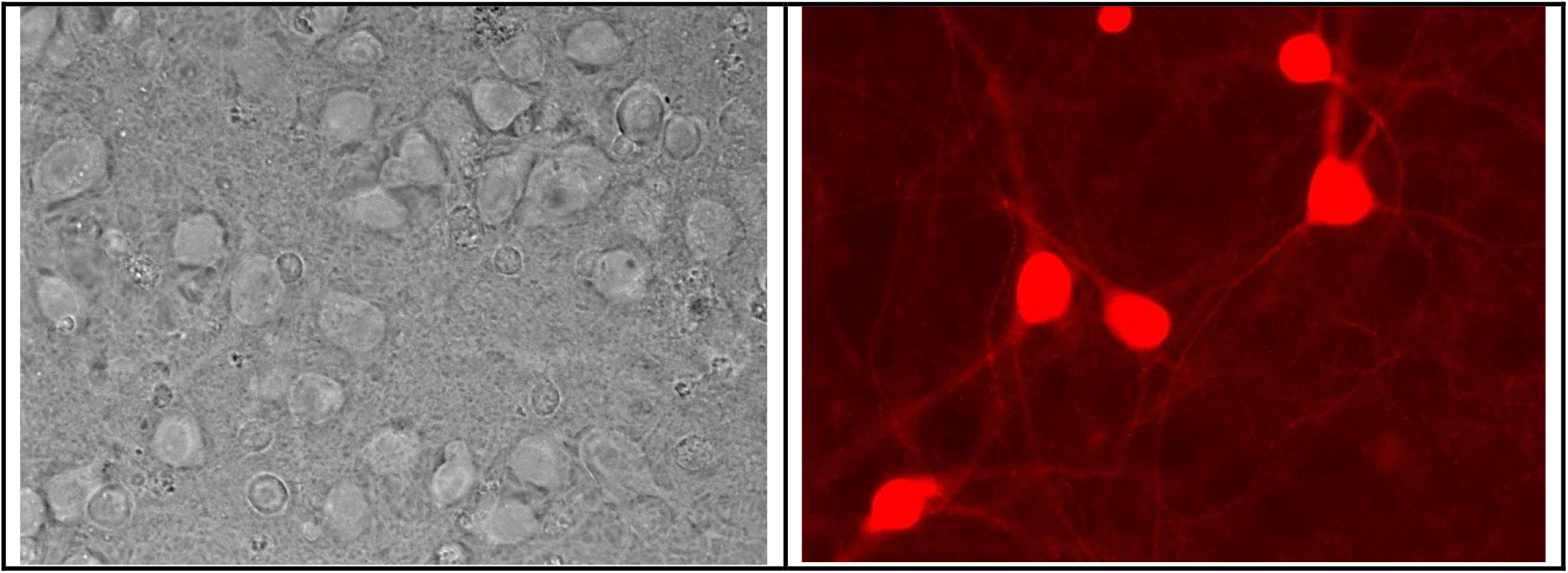
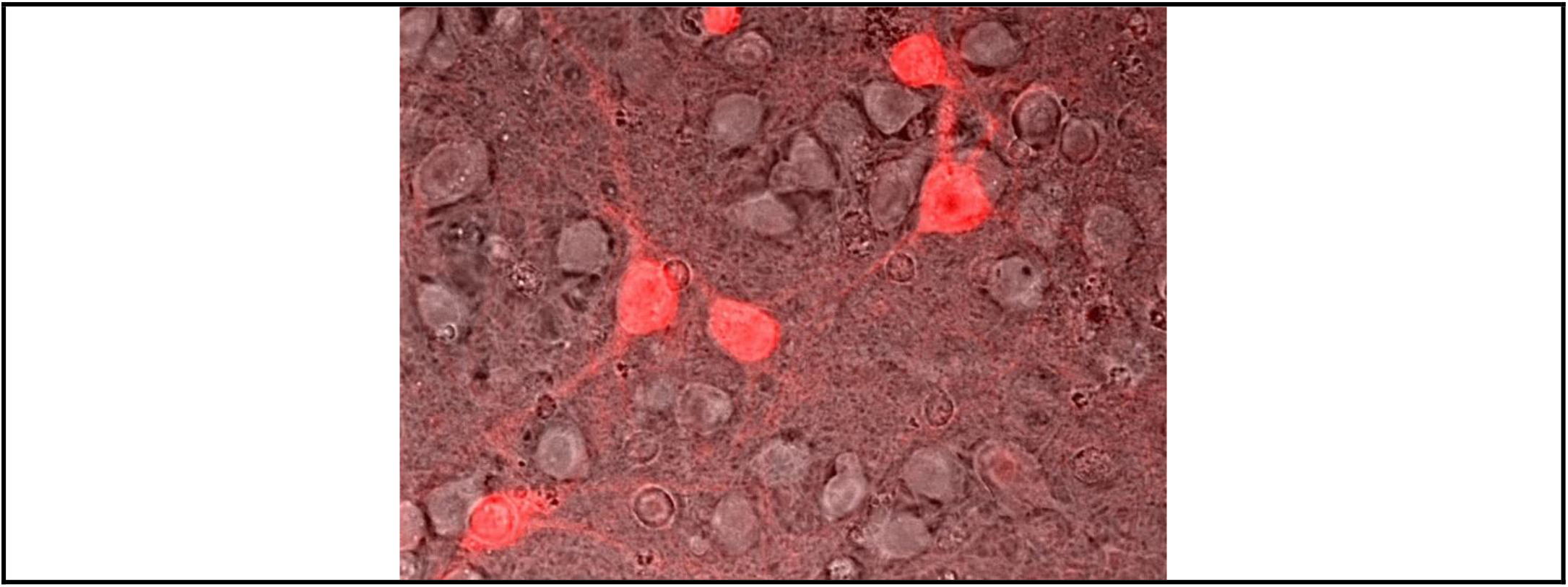
Images of cultured rat neocortical neurons at DIV 19 and transformed with AAV-mDlx-NLS-mRuby2 (40x magnification). (Top Left) Brightfield image of neurons. (Top Right) Fluorescence of mRuby2 in red. (Bottom) Overlay of both images to show the neocortical interneurons in the field.

**Figure 2.**
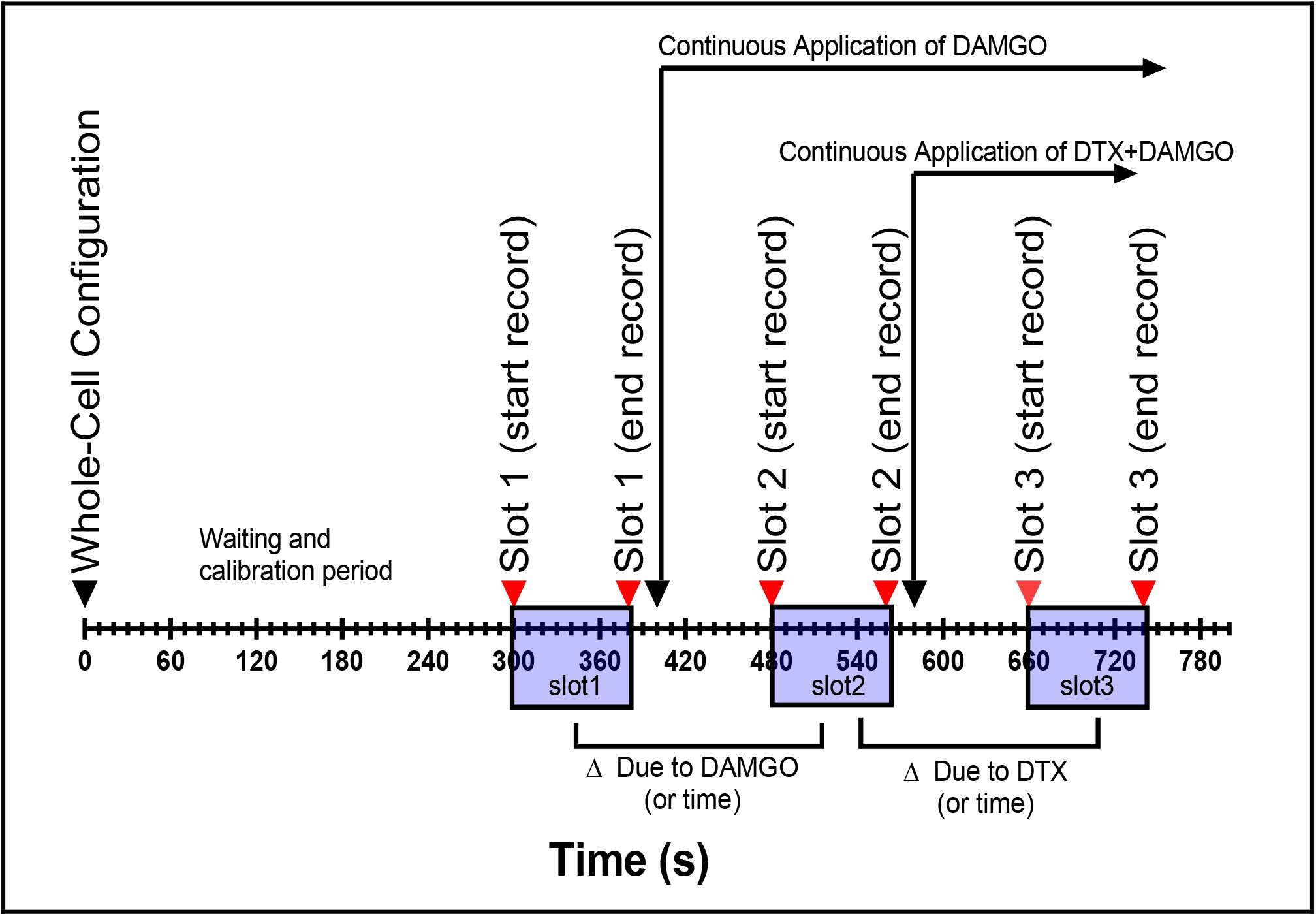
Time course for patch-clamp recordings. Upon achieving whole-cell configuration, we calibrated the amplitude of the current injections to that particular neuron (to factor in its RMP and input resistance) during the waiting and calibration period. (20 μM) CNQX and (50 μM) D-AP5 were constantly perfused to block glutamatergic receptors. We performed the “Initial” (slot 1) recording after this period and applied (3 μM) DAMGO for 80s of incubation and perfusion. We then performed the “post-DAMGO” recording (slot 2) and applied (100 nM) αDTX+DAMGO for another 80s of incubation. After that, we finally collected the “post-αDTX+DAMGO” (slot 3) recording. To evaluate for DAMGO effects, we compared slot 2 with slot 1. To look for αDTX effects in background of DAMGO, we examined the change between slot 3 and slot 2. We also collected recordings from neurons under a saline-control condition to account for the effects of time in the drug group. We recorded from these neurons in an identical fashion (the timeline is the same), but the saline-control neurons only received vehicle buffer, and not DAMGO or αDTX.

We tested for a significant Slot*Group interaction in the H-responders versus the controls in timeslots 2 to 3 (DAMGO → αDTX+DAMGO). This interaction was significant indicating a potential αDTX effect on the 7 variables (*F*(7,19)= 2.57, *p* = 0.048, *Wilks’ λ* = 0.486). We therefore conducted further analysis on each measure. We had previously found significant DAMGO effects in all the measures for H-responders, except for AP threshold and afterhyperpolarization amplitude (see Table 3), but we expected that αDTX’s effects may not be as extensive as DAMGO’s effect, since this channel may only mediate some of the μOR’s inhibitory mechanisms.

We found no significant effect (*t*(20) = 0.33, *p* = 0.374) of αDTX on the RMP of H-responders (*M* = 0.98, *SD* = 0.03) when compared to saline-controls (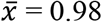, *SD* = 0.04). This indicates that although DAMGO had hyperpolarized these neurons, αDTX-sensitive channels were not responsible for this effect. However, we did find a significant (*U* = 32, *p* = 0.033) effect of αDTX on the AP V_m_ threshold of H-responders (Mdn = 1.04) when compared to the saline controls (Mdn = 1.00). This was a particularly interesting finding because DAMGO did not modulate AP threshold (Figure 6 and Table 3).

Additionally, αDTX significantly reduced ISI (*t*(20) = 2.70, *p* = 0.007) in H-responders (*M* = 0.86, *SD* = 0.13)) versus the saline controls (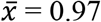, s = 0.04)). Similarly, αDTX significantly increased (*U* = 22, *p* = 0.005) the number of APs in the H-responders (Mdn = 1.22) versus the saline controls (Mdn = 1.00). Therefore, αDTX has an opposite effect to DAMGO, which raises ISI and decreases evoked APs in these H-responders. This suggests that αDTX-sensitive currents do indeed mediate some of the DAMGO effect on suppressing the initiation of APs at suprathreshold depolarizations. In a later analysis, we analyzed whether αDTX fully or partially reverted these properties on a cell-by-cell basis (Figure 9).

We next investigated whether αDTX reversed the DAMGO effects on AP kinetics through halfwidth and maximum rates of repolarization of an AP. αDendrotoxin failed to reverse (*U* = 48, p = 0.219) DAMGO effects in AP halfwidth in the H-responders (Mdn = 1.01) versus the saline controls (Mdn = 1.06). Similarly, we found that αDTX did not significantly (*U* = 57, *p* = 0.423) alter the maximum rate of AP repolarization (Mdn = 0.91) when the H-responders were compared to the saline controls (Mdn = 0.96). Thus, it appeared that the changes in AP waveform induced by DAMGO are mediated by channels other than the αDTX-sensitive channels.

Finally, we investigated whether αDTX altered AHPs. We found that αDTX produced a significant reduction (*U* = 29, *p* = 0.020) in AHP amplitude when H-responders (Mdn = 0.84) were compared to the saline controls (Mdn = 0.98). Therefore, DAMGO increased AHP amplitude, while αDTX reduced it.

### αDTX and magnitude of reversal

In H-responders, αDTX affected ISI, number of evoked APs, AHP amplitude, and AP threshold (However DAMGO had not changed AP threshold). We next wanted to determine whether αDTX was fully reversing the effects of DAMGO in these parameters, or only partially reversing them.

To visualize this variability effects of αDTX in a background of DAMGO, we baselined each of the H-responders to their initial starting point and graphed all 7 changes over the course of slots 1-3 (Figure 9). These analyses provided insight into the direction of the αDTX effect, while also charting each neuron’s course individually. From these graphs it is evident that not only did the effect of DAMGO and αDTX varies between the interneurons, which can be expected of a heterogenous population of neocortical interneurons.

### DAMGO alters RMP and AP waveform in S-responders

We began testing for DAMGO effects in S-responders by testing the 7 combined variables in a mixed-model repeated measures MANOVA for a significant Slot*Group interaction. We found a significant effect of DAMGO in S-responders (*F*(7,20) = 2.57, *p* = 0.046, *Wilk’s λ* = 0.473) and we therefore investigated them further.

We first compared their change in RMP (*M* = 1.02, *SD* = 0.04) against saline controls (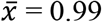, *s* = 0.02). Apparently, although this group fell short of the threshold for the H-responder group compared to the saline group (Figure 3) the S-responders too had undergone a significant hyperpolarization (*t*(18) = 2.53, *p* = 0.011) compared to the saline controls. Although these S-responders underwent statistically significant hyperpolarization after DAMGO, we classify them as S-responders because they were not selected by that particular criterion, as the H-responders had been.

**Figure 3.**
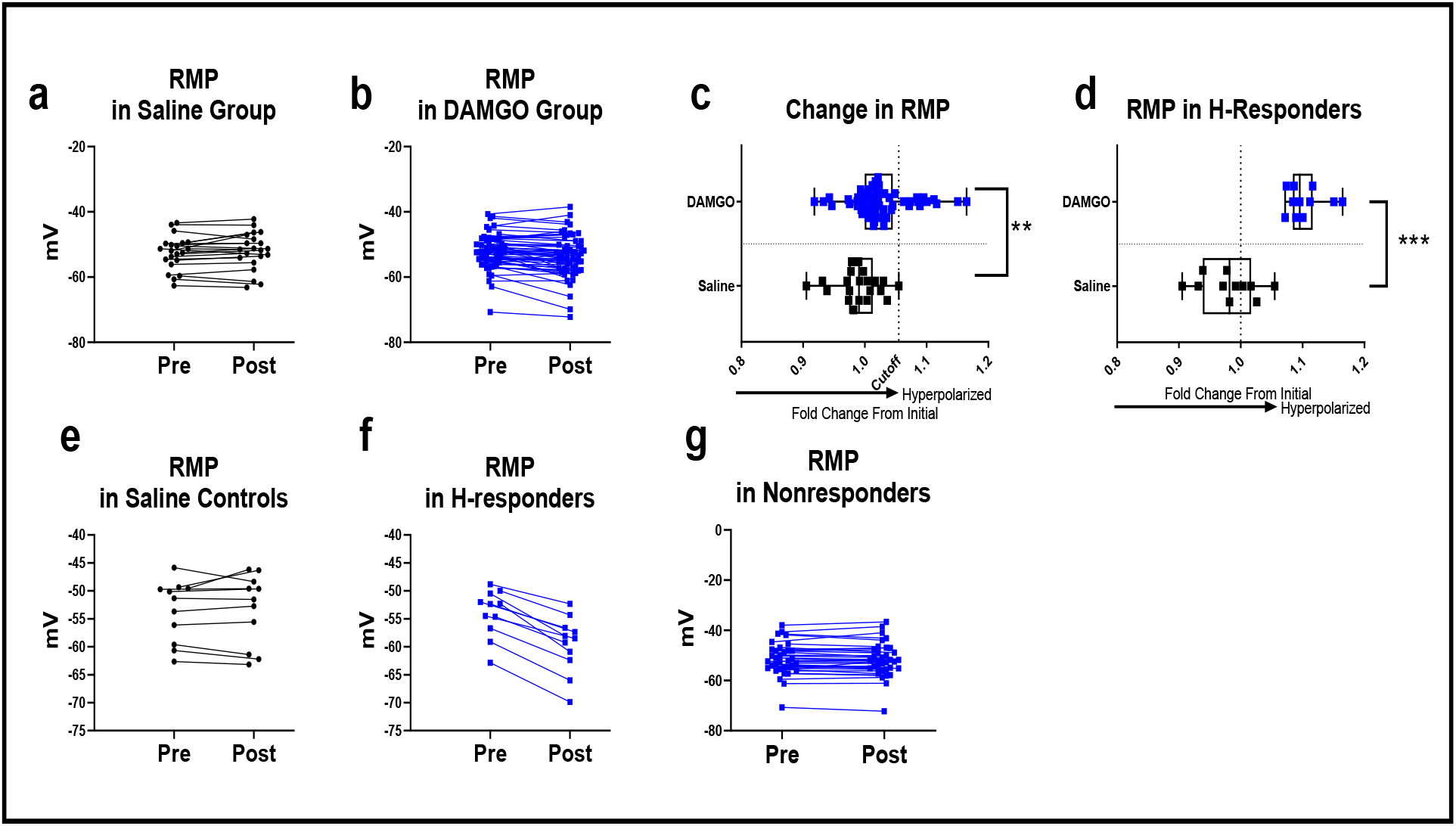
H-responders hyperpolarize. The efforts to identify DAMGO-responders and measure the change of RMP in the H-responders began by graphing resting membrane potential (RMP) after the 80s period of incubation with either (3 μM) DAMGO or vehicle saline (a) We plotted the RMP each saline-control neuron (*N* = 21) individually and (b) each DAMGO-exposed neuron (*N* = 55) individually. However, these graphs were ineffective in identifying the neurons that hyperpolarized the most under the DAMGO condition. To isolate the change (hyperpolarization), we graphed the fold change in RMP instead of the raw value (c) We plotted the change in RMP (defined as slot 2 RMP/ slot 1 RMP) in saline and DAMGO controls and found a significant hyperpolarization in the DAMGO group compared to the saline group (*p* < 0.05). To identify true DAMGO responders, we established a cutoff at the most hyperpolarized saline control (vertical dotted line). DAMGO hyperpolarizers (to the right of the cutoff) are hereafter called “H-responders” (*N* = 11). (d) We extracted these H-responders and a comparison group of random saline controls (n = 11) onto a new plot. This represents the change in RMP among the H-responders, which was significantly (*p* < 0.05) more hyperpolarized than the random controls. (e) Subset of random saline controls for comparison (*n* = 11) showing little general trend towards depolarization or hyperpolarization during the slot1 to slot 2 period. (f) A plot of the H-responders (*N* = 11) RMP in their actual values showing a trend towards hyperpolarization. (g) The RMP of the nonresponders (*N* = 34) during the same period. (a,b,e,f,g) Dots are values for interneurons and lines connect the same neurons before and after saline or the DAMGO (c,d) Edges of box are 1^st^ and 3^rd^ quartiles for values, dots are the change in RMP for the individual neurons, and whiskers extend to the largest and smallest values. Statistical testing performed with unpaired t tests (one-tailed) after confirming their data had a normal distribution with a D’Agostino-Pearson omnibus K2 test (*p* > 0.05).

**Figure 4.**
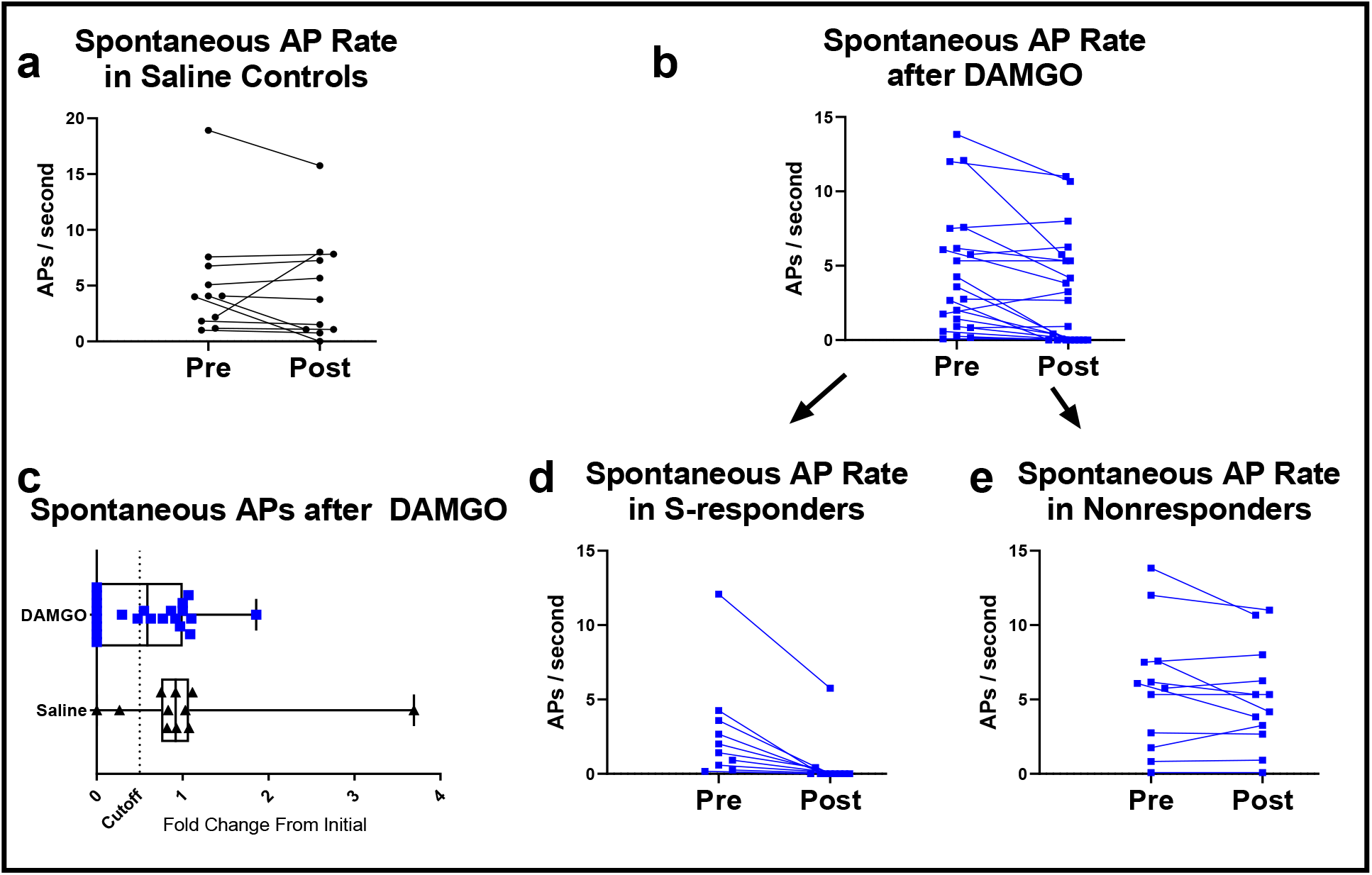
Spontaneous APs after DAMGO or saline. Out of the remaining pool of 44 (3 μM) DAMGO-exposed neurons, 22 were spiking spontaneously. Out of the 21 saline controls, 11 were spiking spontaneously. (a) Rate of spontaneous APs after 80s of vehicle saline show no general trend. (b) Rate of spontaneous APs in the remaining (non-H-responder) DAMGO-exposed neurons, which appeared to show some neurons had reductions in spontaneous APs, and others had not. (c) We therefore graphed the fold change in spontaneous APs after DAMGO or vehicle saline. We established a 50% reduction (vertical dotted line) in spontaneous APs as a somewhat arbitrary cutoff to distinguish nonresponders from S-responders. (d) Rate of spontaneous APs in S-responders (*N* = 10) which had fell below the cutoff, and therefore were classified as S-responders. (e) Rate of spontaneous APs in the remaining pool of nonresponders (*N* = 12), which had not achieved that 50% cutoff. Figures (d) and (e) are therefore subsets of (b).

**Figure 5.**
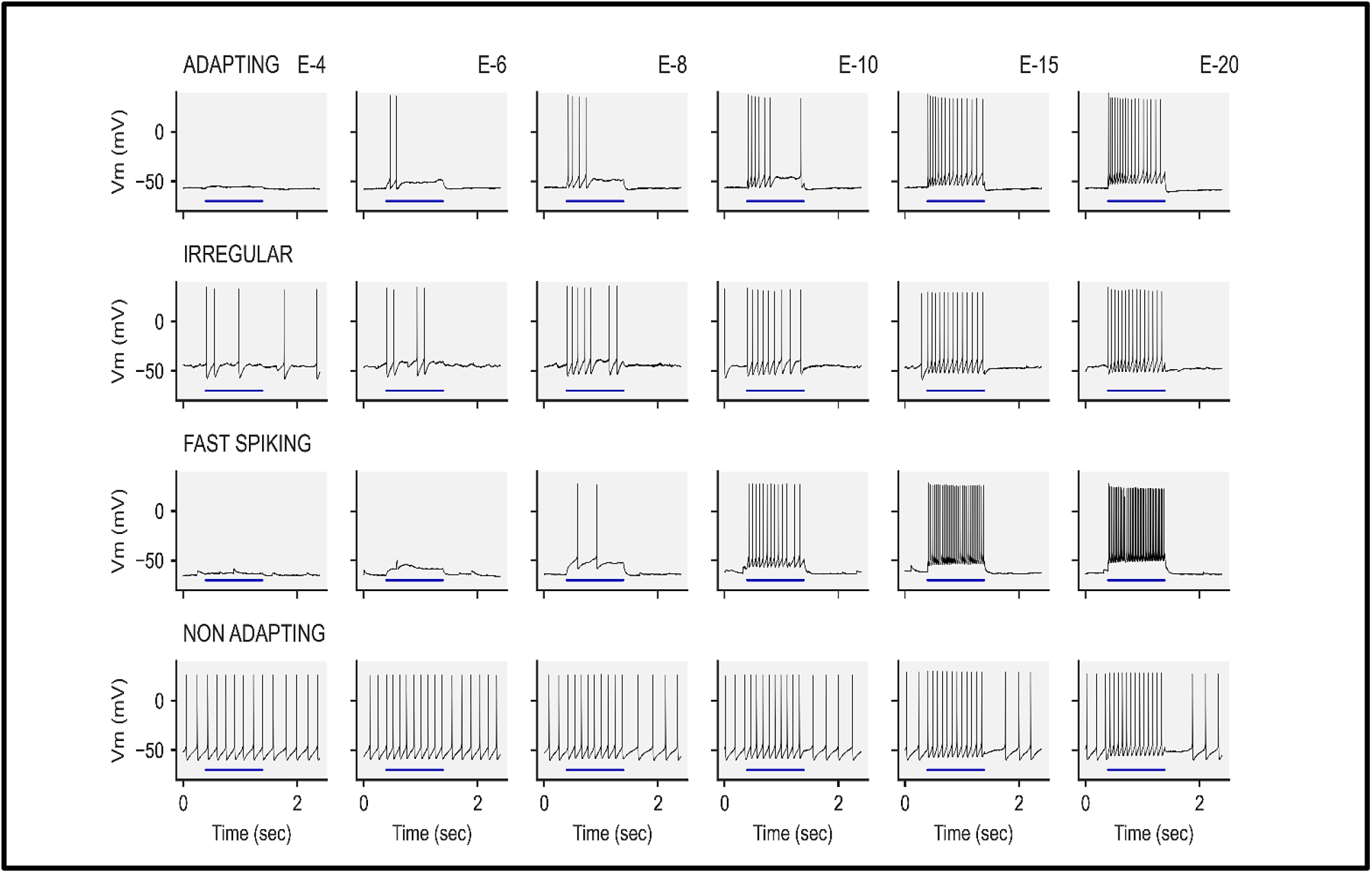
Representative Examples of Firing Patterns. We classified the spiking characteristics of interneurons by observing the pattern of AP discharges that occurred around their V_m_ threshold for an AP. Note that there is some variability here in their threshold and RMP, which manifests as threshold V_m_ sometimes occurring at different episodes for each neuron. Firing patterns are arranged by row, and their episodes by column. (Top row) Adapting neurons tended to fire more frequently at the beginning of the current pulses and had steadily-increasing interspike intervals through the current application. This pattern stayed stable even at high current applications (E-15 and E-20). (Second Row) Irregular spiking neurons fired irregular APs, or irregular bursts of APs, at suprathreshold current application. However, this firing pattern often transitioned to more evenly spaced APs at higher current applications. (Third row) Fast-spiking neurons tended to have more distance between their RMP and AP threshold. Most of their APs were evenly spaced throughout the current application. At higher current applications (E-15 and E-20) this firing pattern tended to fire more frequently, and correlate with the amplitude of the current injection. (Fourth Row) Nonadapting, nonfastspiking neurons had evenly spaced spikes throughout the current application. This firing pattern tended to have a stable discharge pattern regardless of the current application; unlike the neurons we classified as fastspiking which tended to increase their AP frequency when more-depolarizing currents were applied.

**Figure 6.**
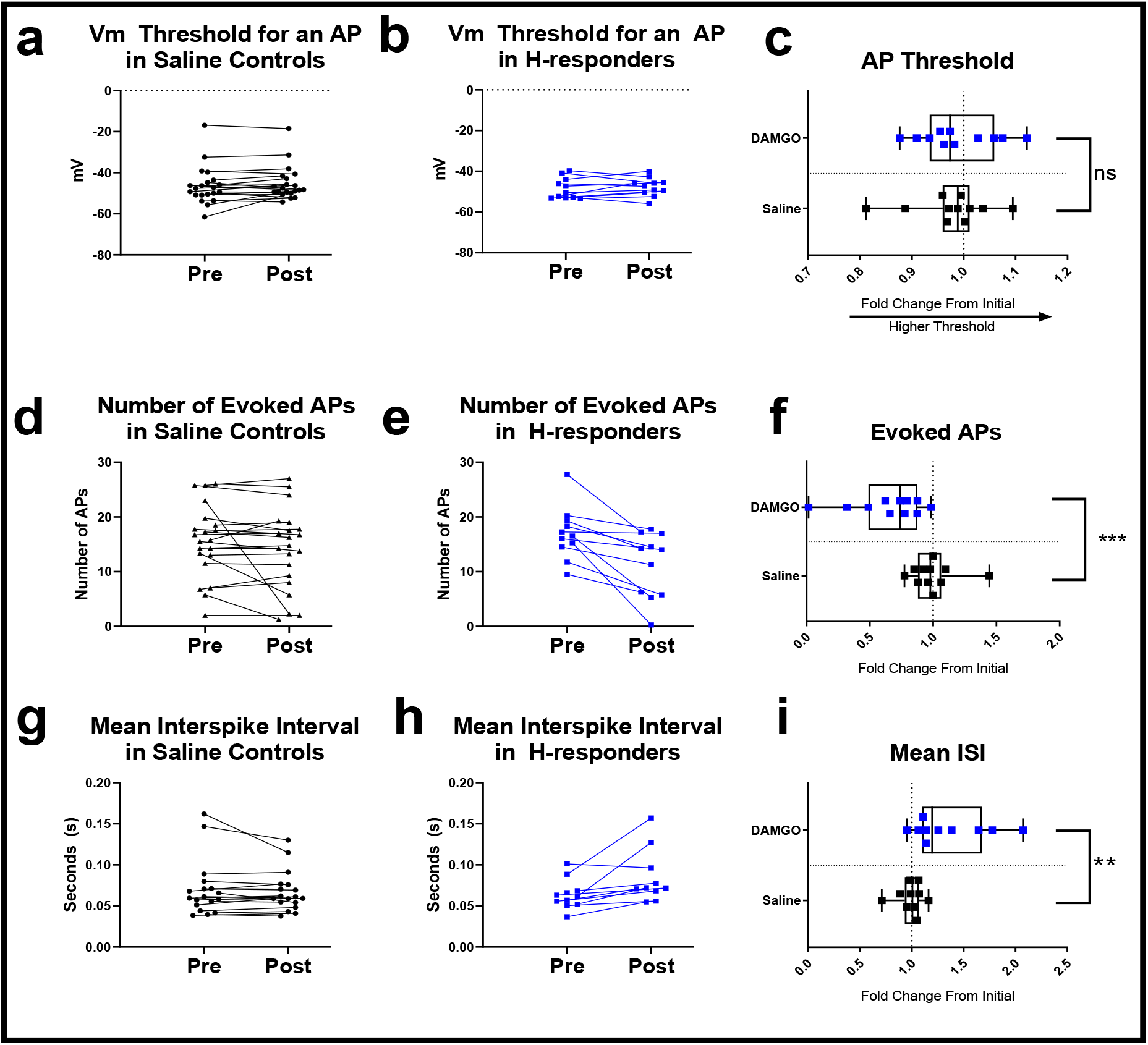
Spike frequency altered in H-responders. We tested whether H-responders had discernible changes after (3 μM) DAMGO AP threshold and frequency by comparing them to saline controls. (Left column) The saline controls (*N* = 21) show little systematic changes in their (a) AP threshold, (d) number of evoked APs, (g) and mean interspike interval during exposure to vehicle saline. (Middle column) Meanwhile the H-responders (*N* = 11) show little change after (3 μM) DAMGO in (b) AP threshold, however they generally show reduced numbers of (e) evoked APs, and (h) larger interspike intervals after DAMGO. (Right column) To illustrate and test these changes more precisely, we measured the fold change in those parameters against a random group of saline controls (*n* = 11). We found that (c) AP threshold change was unaffected by the DAMGO compared to the change in saline controls, but we found significantly-decreased (*p* < 0.05) number of (f) evoked APs in H-responders after DAMGO and significantly increased (i) mean interspike interval. One H-responder is excluded because it became single-spiking after DAMGO (Left and Middle columns) connecting lines bridge the same neuron after an 80s period of vehicle or DAMGO incubation. Y axes are true values for those particular measures (Right column) isolates the fold change (slot 2/slot1; postdrug/initial) after vehicle saline or DAMGO. Edges of the boxes are drawn at the 1^st^ and 3^rd^ quartiles while the “whiskers” connect the largest and smallest values for that group. We performed statistical testing of ISI and AP threshold with unpaired t-tests (one-tailed) after first testing for skewness and kurtosis with D’Agostino-Pearson omnibus K2 test (*p* > 0.05). Only number of evoked APs was significantly different from a normal distribution (*p* < 0.05) and therefore tested with a one-tailed Mann-Whitney U test.

**Figure 7.**
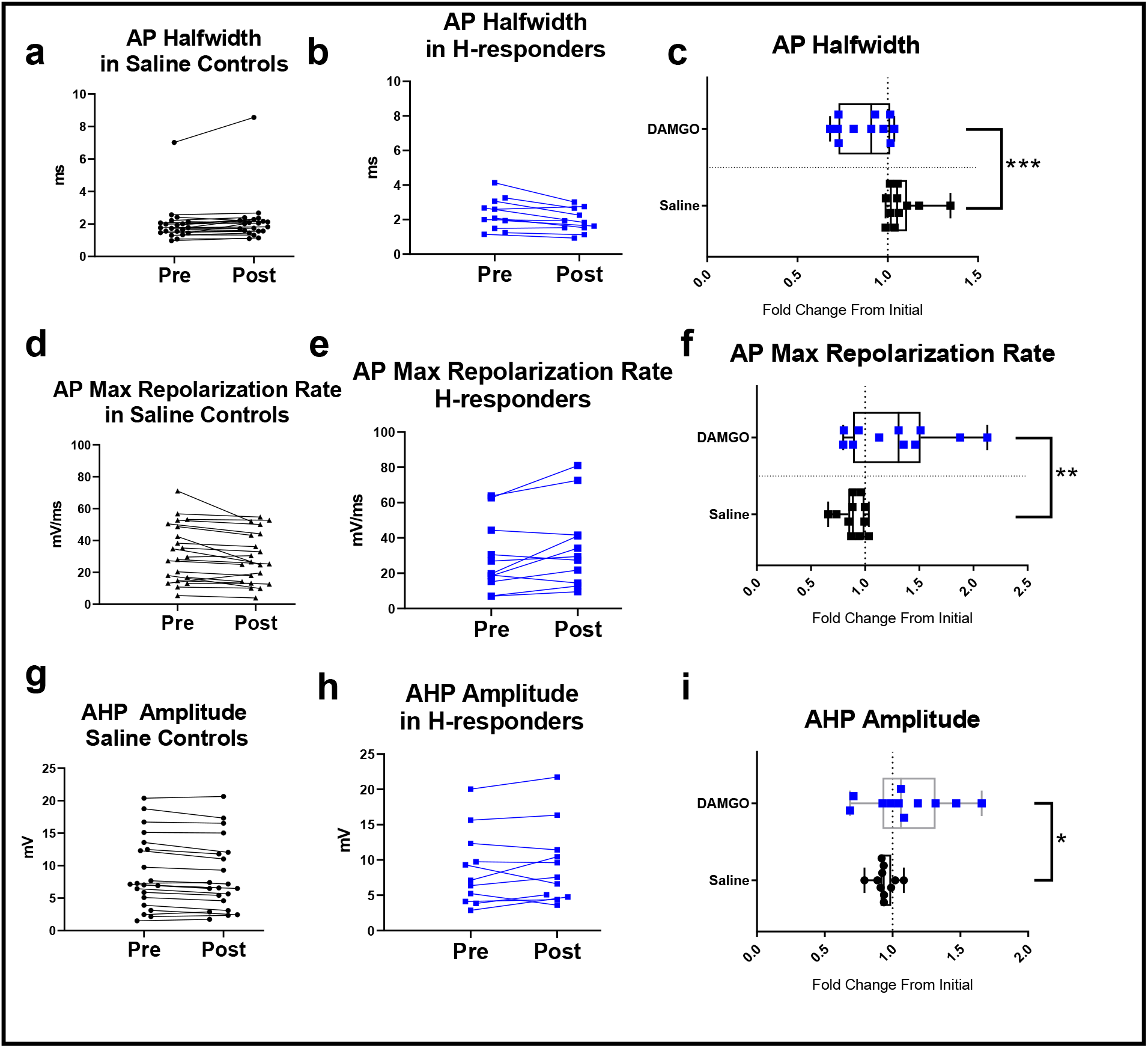
AP kinetics are altered in H-responders. We analyzed AP kinetics and afterhyperpolarization (AHP) amplitude before and after DAMGO or vehicle saline to determine whether DAMGO altered these properties. (Left column) Saline controls (*N* = 21) before and after an 80s period of exposure to the vehicle saline. In the saline group, there was little trends in (a) AP halfwidth, and (d) maximum repolarization rates (Middle column) DAMGO-exposed H-responders (*N* = 11) generally show reduced (b) halfwidths and (e) larger maximum repolarization rates after DAMGO, but (h) AHP amplitudes are generally stable. (Right column) To isolate and statistically test the change after DAMGO, we plotted the fold change after DAMGO or vehicle saline. We found that DAMGO exposure in H-responders resulted in significantly reduced (c) halfwidths compared to random saline controls (*n* = 11). This corresponded with significantly increased (f) maximum AP repolarization rates. We also found significantly larger (i) AHP amplitudes in H-responders relative to controls. (Left and Middle columns) connecting lines bridge the same neuron after an 80s period of vehicle or DAMGO incubation. Right column isolates the fold change (slot 2/slot1; postdrug/initial) after vehicle saline or DAMGO. Edges of the boxes are drawn at the 1^st^ and 3^rd^ quartiles while the “whiskers” connect the largest and smallest values for that group. We performed statistical testing for maximum repolarization rate and AHP amplitude with unpaired t-tests (one-tailed) after testing for skewness and kurtosis with D’Agostino-Pearson omnibus K2 test. AP halfwidth change (for saline group) was nonnormally distributed (*p* < 0.05) and therefore tested with Mann-Whitney U tests.

**Figure 8.**
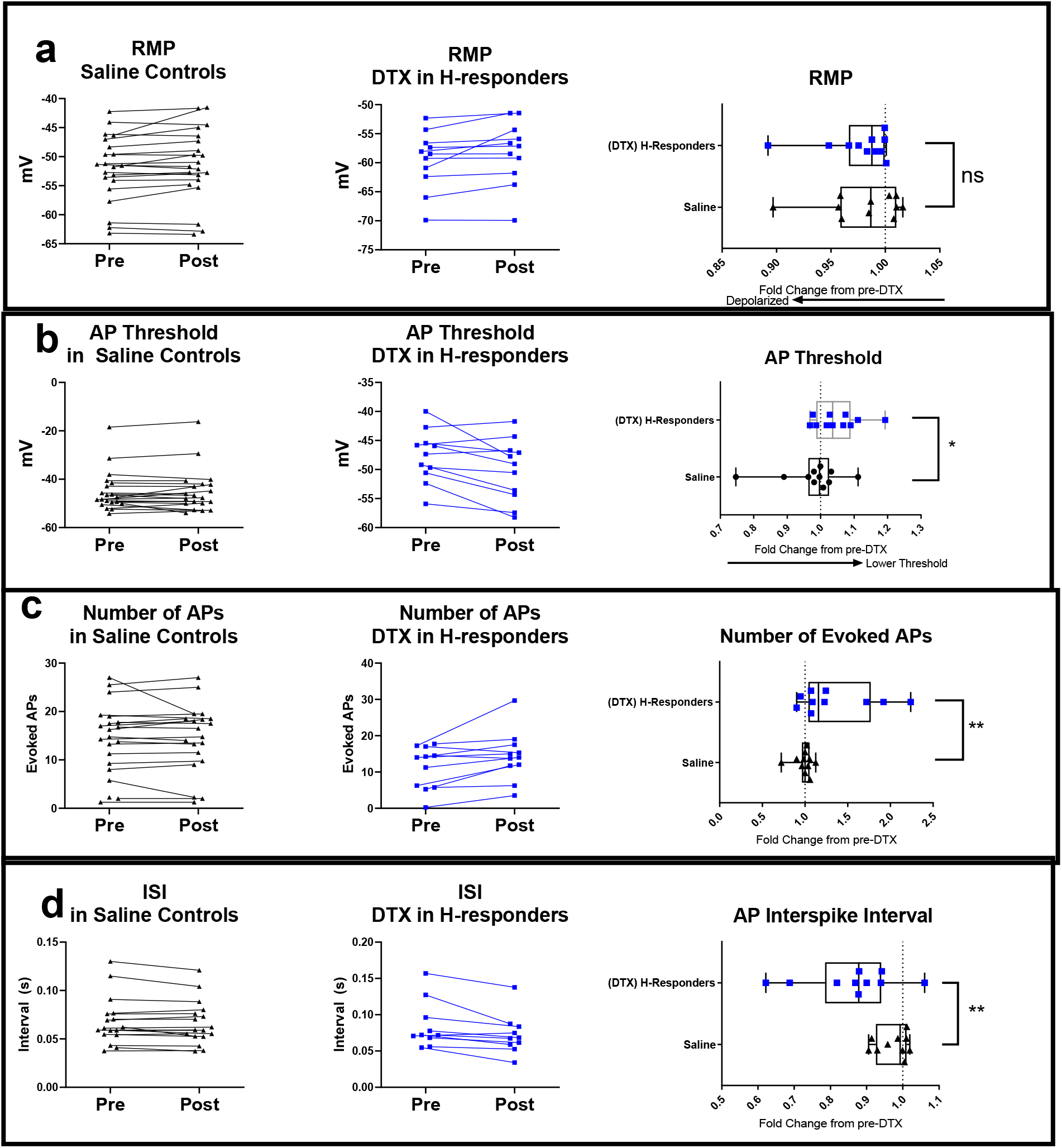
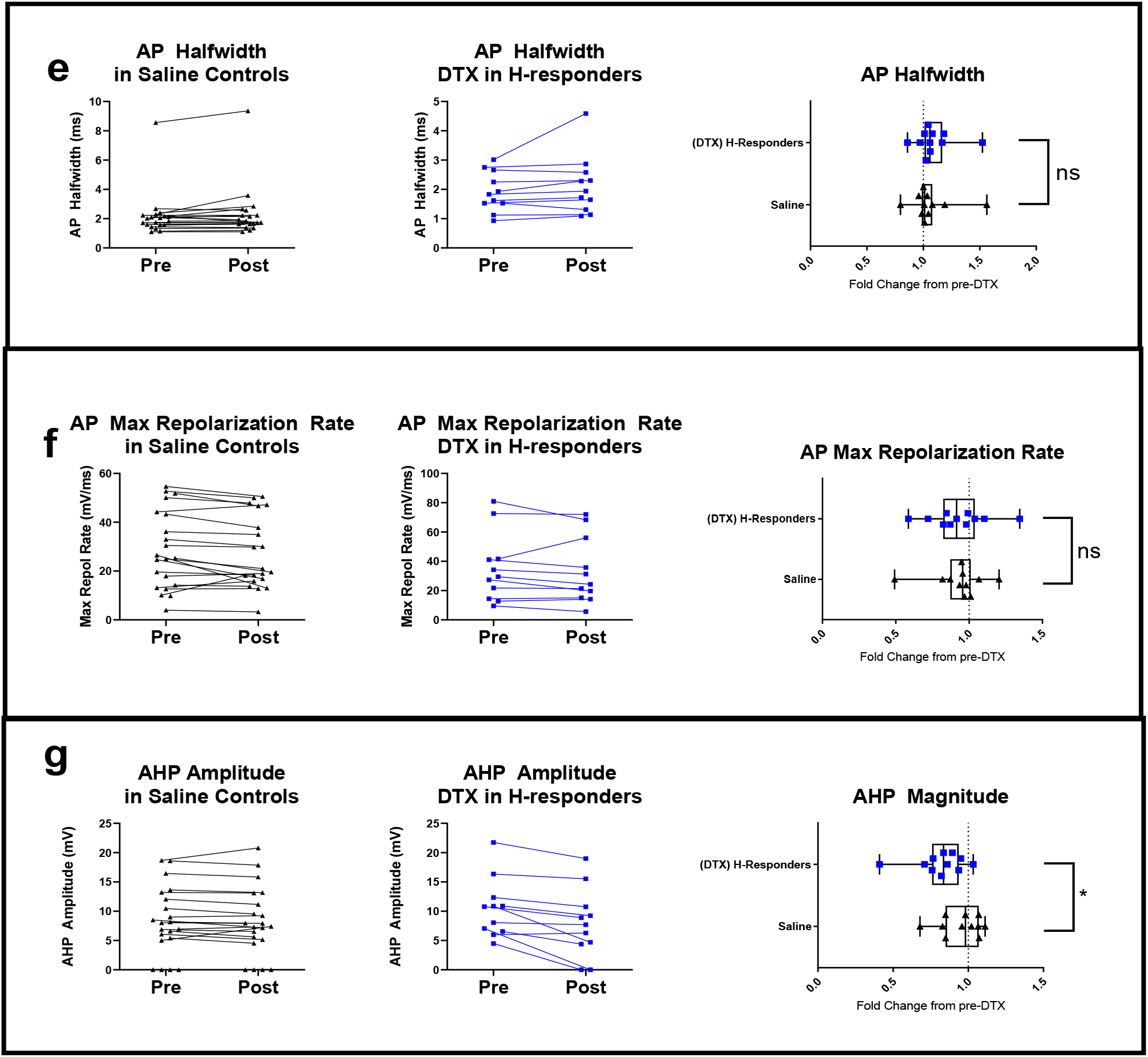
αDTX reverses DAMGO effects on ISI and AP number. We tested for αDTX effects in the H-responders by applying αDTX in a background of DAMGO and examined these parameters for changes that were in opposite polarity as DAMGO. These change ratios were calculated by dividing postDTX/preDTX (i.e., slot 3/slot 2). Thus, αDTX in a background of DAMGO. (Left column) Real values for saline controls during this period which show stochastic changes in each measure. (Middle column) Real values for H-responders before and after exposure to αDTX. (Right column) isolates the fold change in a subset (*n* = 11) of saline controls and the (*N* = 11) H-responders from their pre-DTX condition. Statistical testing was performed by comparing the changes in saline controls (*n* = 11) with the changes in H-responders (*N* = 11) after αDTX. We found no significant effects of αDTX on (a) RMP, (e,f) AP kinetics (halfwidth and max repolarization rate). However, we did find significant effects of αDTX on H-responders in a background of DAMGO in (d) interspike interval (c) number of evoked APs, as well as (g) afterhyperpolarization amplitude. We also found that αDTX lowered (b) AP threshold, which had not been modulated by DAMGO. One neuron was excluded from graph (c), but not statistical comparison, due to a 14-fold increase in evoked APs. Edges of the boxes are drawn at the 1^st^ and 3^rd^ quartiles while the “whiskers” connect the largest and smallest values for that group. We performed statistical testing with unpaired t-tests (one-tailed) after testing for skewness and kurtosis with D’Agostino-Pearson omnibus K2 test (*p* > 0.05).

**Figure 9.**
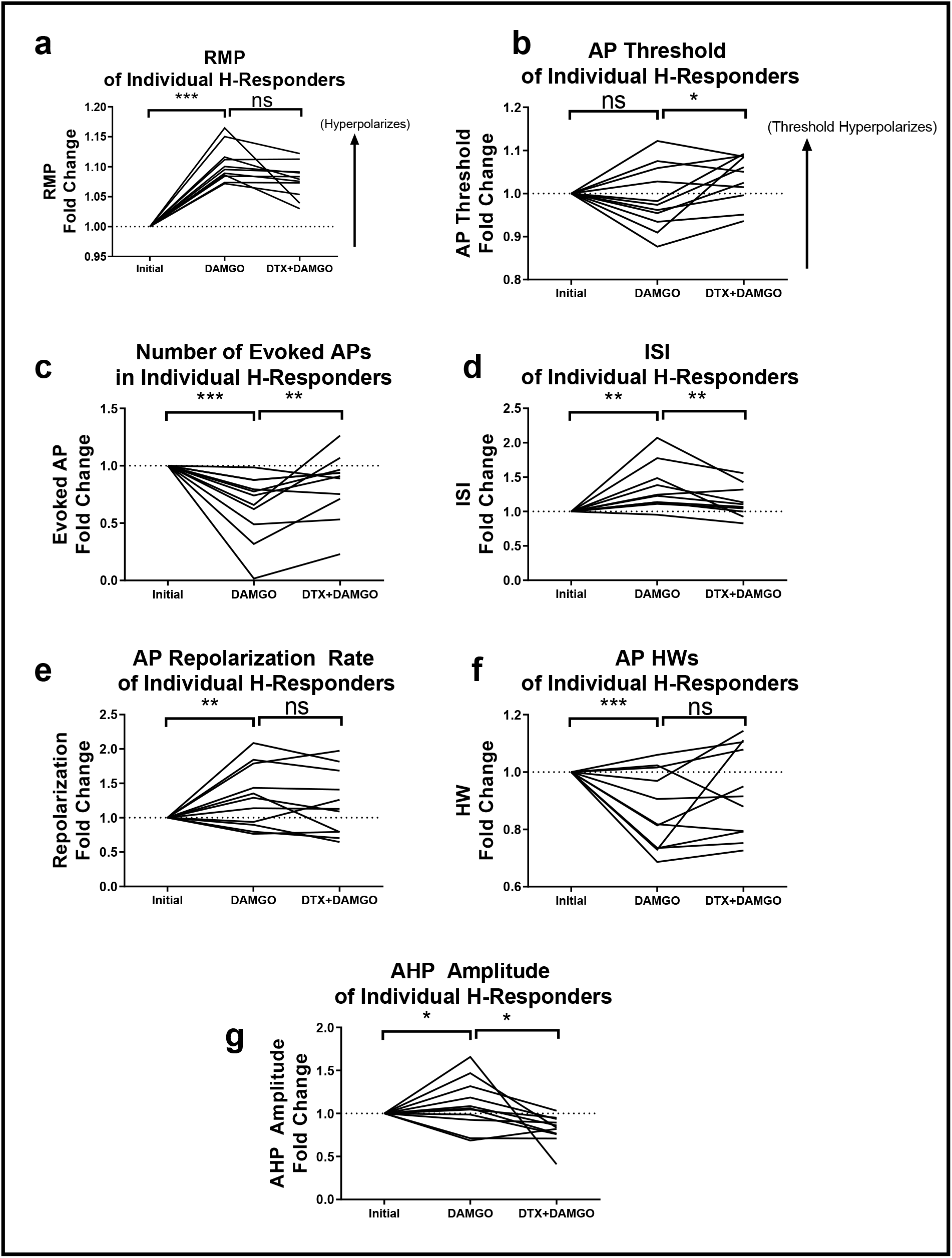
αDTX partially reverses some effects of DAMGO in H-responders. To illustrate the fold changes in all 7 parameters across the 3 timepoints, we baselined value for these measures in the (*N* = 11) H-responders by their initial value; therefore, all neurons emanate from “1.00” which marks their original values. The dotted line at 1.00 represents baselined starting value for all neurons. These graphs show the fold changes over the course of their treatment with DAMGO, and then αDTX+DAMGO. (a) Change in RMP, which was the basis of selection for (*N* = 11) H-responders, and not significantly reversed by αDTX. (b) Change in AP threshold, which was not significantly altered by DAMGO, shifted towards hyperpolarized V_m_ after αDTX. (c) Evoked APs in H-responders and (b) ISI changes in H-responders (*n* = 10; one neuron was not included because it became single-spiking after DAMGO). Changes in (c,d) had before been found to be affected by both DAMGO and αDTX in H-responders. (e) Maximum repolarization rate and (f) AP halfwidth, which were both significantly changed by DAMGO, but not αDTX. (g) Afterhyperpolarizations were significantly enhanced by DAMGO and reduced by subsequent addition of αDTX. Lines on all graphs connect the same neuron’s values across all 3 timepoints. Statistical comparisons were made by comparing fold change in H-responders to fold changes in saline controls. These statistical comparisons were displayed previously and consolidated into these graphs for convenience (Figures 6, 7, and 8; Tables 3 and 5). Changes were compared for significance with unpaired t-tests or Mann-Whitney U tests.

We tested their fold change in AP threshold to find a nonsignificant effect of DAMGO (*t*(18) = 1.11, *p* = 0.141) when S-responders (*M* = 0.96, *SD* = 0.08) versus the saline controls (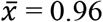, s = 0.07); there was no effect of DAMGO on V_m_ threshold for an AP in the S or H-responders.

We did not find a significant difference in their changes in interspike interval (*t*(18) = 1.42, *p* = 0.086) of these S-responders (*M* = 1.09, *SD* = 0.12) relative to the saline controls (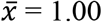, *s* = 0.14). Nor did we find a significant (*U* = 30, *p* = 0.072) decrease in number of evoked action potentials (Mdn = 0.90) relative to the saline-only controls (Mdn = 1.01). Interestingly, we found effects in these 2 properties in the H-responder group but apparently these effects are absent in the S-responder category; although these neurons were selected for on the basis of their decreased spontaneous APs, it appears that somatically-evoked APs were not similarly decreased or reduced in frequency.

We next analyzed the AP kinetics of the S-responders to determine whether they too had altered AP shapes after DAMGO as the H-responders did. We found that the S-responders had significantly (*t*(18) = 2.42, *p* = 0.013) reduced their AP halfwidths (*M* = 0.95, *SD* = 0.13) after DAMGO compared to saline controls (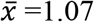, *s* = 0.10). These S-responders (Mdn = 1.07) also had a significant increase (*U* = 19, *p* = 0.009) in their maximum rate of repolarization when compared with saline controls (Mdn = 0.87).

Finally, we analyzed the fold change in afterhyperpolarizations in S-responders (*M* = 1.07, *SD* = 0.24) versus changes in the saline control (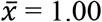, *s* = 0.09). However, there were no significant differences in the changes of their AHP amplitude (*t*(18) = 1.72, *p* = 0.051).

To summarize, spontaneous AP responders had relatively small hyperpolarizations, and changes in AP kinetics compared to H-responders (Table 7). S-responders lacked the DAMGO-induced ISI increase and reduced evoked AP number, which were observed in H-responders (Table 3). Thus, it appears that DAMGO influenced their RMP, spontaneous APs frequency, and AP repolarization kinetics, but it did not change the somatically-evoked AP frequency, as DAMGO had done in the H-responders. The effects of DAMGO in S-responders appeared to be smaller in magnitude and in the range of affected parameters (Table 7)

### αDTX effects absent in S-responders

We also investigated whether there were αDTX effects in the S-responders, although we did not expect a αDTX effect here since data from the H-responder suggested that the effects of αDTX were restricted to AP threshold, ISI and number of evoked APs, none of which were affected by DAMGO in the S-responders. We found a non-significant Slot*Group interaction (*F*(7,19) = 0.81, *p* = 0.592, *Wilks’ λ* = 0.229) in this group for the combined variables, indicating that there was not an effect of αDTX in the S-responders.

We also wanted to determine whether αDTX was reversing the measurement that was used to initially identify the S-responders; their spontaneous APs. However, the number of spontaneous APs was not significantly changed when αDTX was applied in a background of DAMGO (*U* = 32, *p* = 0.069). Indeed, after DAMGO exposure, only 3 out of 10 S-responders were firing spontaneously, and only 1 out of 7 had resumed firing spontaneous APs after αDTX. Thus, it appears that DAMGO-induced downregulation of spontaneous APs was not detectably related to αDTX-sensitive channels.

### Investigating the influence of RMP polarization

The identification of H-responders to DAMGO involved the selection of neurons with the largest hyperpolarization in during those 80s of DAMGO incubation. Under that procedure, it was possible that any subsequent differences we observed was simply a consequence of that difference in polarity change between the sampling periods. To investigate the confound of polarization, we selected the neurons that had hyperpolarized the most in the saline control, and we also selected a comparison group of neurons that had depolarized the most in the DAMGO group. Although we expected that this depolarizing change was merely due to chance (and not a DAMGO effect) it enabled us to “flip the sign” so that the saline-control neurons were relatively more hyperpolarized than the DAMGO-exposed neurons. If all of these changes were actually a result of hyperpolarization, the most-hyperpolarizing saline neurons should actually undergo analogous changes as the H-responders. If they fail to do this, significant effects in AP parameters are more likely to be caused by differential ion channel regulation by μORs, rather than a polarizing shift in the V_m_ without ion channel regulation by μORs.

We found a significant Slot*Group interaction (*F*(7,12) = 2.57, *p* = 0.048, *Wilks’ λ* = 0.633) in these groups for the combined dependent variables, which included the RMP change that they had been selected on. We therefore investigated them further for changes in other parameters.

Other than the change in RMP that they had been selected on, we did not find significant changes in any of the 6 other parameters. These analyses suggest that parameters following RMP were not being influenced by the change in the magnitude of their polarity that served as the maker for H-responders, but instead were a result of differential ion channel regulation that occurs during a DAMGO-response.

### Investigating false negatives

We next wanted to determine whether there were DAMGO-induced changes in the nonresponder group, which would have suggested that these groupings were too exclusionary. This could contribute false negatives, i.e., true responders that were excluded from the responder groups and passed unanalyzed into the pool of “nonresponders.” Out of the 55 neurons that we recorded and exposed to DAMGO, there remained 34 recordings that were excluded from the S&H-responder categories. We wanted to investigate this nonresponder pool for the same changes in AP parameters that we sought in the S&H-responder categories. We decided to begin to address this issue by seeking changes in the nonresponders for the 7 measures we have been testing for.

We individually graphed the change in polarity (x axis) versus all 7 measures tested (y axis). If neurons had undergone changes in those measures without hyperpolarization, they can be visually identified by appearing vertically-displaced, but horizontally close to the graph’s origin (Figure 14).

This comparison revealed that these 2 categories of responders had captured most of the large changes in those parameters, and yet there seemed to be a small cluster of “nonresponders” that seemingly had hyperpolarized and underwent very small reductions in halfwidth and small increases in AP repolarization rates. For instance, the saline-controls never fall in the hyperpolarized and quickly-repolarizing top-right quadrant (Figure 14j) and yet 7 nonresponders do. Therefore, examining for hyperpolarization seemed to predict the larger changes in these measures, but it left open the possibility that weaker responders were being misclassified as DAMGO-nonresponders.

To investigate false negatives that were overlooked, we tested for a significant Slot*Group interaction in the nonresponders with a mixed model repeated measures MANOVA (*N* = 34) versus the saline controls (*N* = 21). However, the result for the combined variables was nonsignificant (*F*(7,44)= 1.17, *p* = 0.342, *Wilks’ λ* = 0.156) despite the higher sample size when compared to S and H-responders. While it’s possible that some responders with weak effects (Figure 14) may have been misclassified as nonresponders, the neurons with the largest changes in measures that we were testing for seem to have been identified as H or S-responders and thus were previously analyzed here.

### Post-hoc analyses on DAMGO, αDTX and AP kinetics

Finally, we conducted a *post-hoc* analysis of 47 other parameters and ratios that have not been previously addressed here. This included individually analyzing sequential APs to determine if the drugs had effects only one specific APs in their spike train, as well as examining ratios for changes between sequential APs. For these post-hoc analyses, we only considered H-responders for the DAMGO and αDTX effects, because only the H-responders had systematic responses to αDTX and had stronger responses to DAMGO (Table 7).

We found that DAMGO was affecting a very wide range of AP kinetic parameters across many sequential APs, with the notable exception of spike amplitude. However, αDTX’s effects were not seen in these parameters (Table 8). Broadly speaking, it appears that DAMGO consistently alters many different features, from individual AP kinetics, to the probability of firing an AP. Meanwhile αDTX-sensitive channels do not seem to contribute the shape of APs in neocortical interneurons, and its effects appear to be restricted to the probability of firing an AP and their firing frequency.

## Discussion

Previous research indicates that the μOR exerts an excitatory effect on cortical networks by suppressing inhibitory neurons (Homayoun & Moghaddam, 2007; Jiang et al., 2019; H. K. Lee, Dunwiddie, & Hoffer, 1980; Madison & Nicoll, 1988; Pang & Rose, 1989; Zieglgansberger et al., 1979). The experiments here likewise demonstrate that the μOR exerts a strong inhibition on many cortical interneurons. We found that over a third (21/55; 38.1%) of the sample of interneurons either hyperpolarized (H-responder) or had reduced spontaneous action potentials (S-responder). The influence of DAMGO in these two populations of interneurons had overlapping characteristics but were distinct. For instance, in H-responders DAMGO elicited clear hyperpolarization and affected AP kinetics and firing frequency. In contrast, the S-responders exhibited relatively smaller hyperpolarizations and small changes in AP kinetics, and no alterations in the probability of somatically evoked APs.

The high degree of variability of neocortical interneurons between and within their classes makes identification of interneurons in our system difficult. We did not attempt cell-marker identification of the neocortical interneurons since identification would have required a thorough multimodal analysis of their characteristics (The Petilla Interneuron Nomenclature, 2008). Particularly because our system enabled more expeditious data collection at the expense of laminar arrangement - itself an identifying characteristic for interneurons.

We found that most H-responders displayed an Adapting firing pattern, characterized by steadily-increasing interspike intervals. This is noteworthy because this spiking pattern was found at much lower rates in the non-responders and S-responders (Table 2). Other researchers have found interneurons with this firing pattern amongst neuropeptide Y-expressing interneurons (Bruno Cauli et al., 1997; B. Cauli et al., 2004; Férézou et al., 2007; Toledo-Rodriguez, Goodman, Illic, Wu, & Markram, 2005; Yun Wang, Gupta, Toledo-Rodriguez, Wu, & Markram, 2002; Y. Wang et al., 2004). This firing pattern in NPY+ interneurons is particularly common amongst Layer I interneurons of the neocortex (Karagiannis et al., 2009), which others have found to express high rates of μORs and hyperpolarize strongly to DAMGO exposure (Férézou et al., 2007), which was the distinguishing feature of H-responders in our study but found in both groups of responders.

**Table 2.** Responder groups demonstrate different proportions of spike patterns. We used the PING classification system to categorize each group (by hand) based on their spiking pattern in response to a 1s depolarizing current at 2-3 steps above threshold, where the trace stabilizes, and multiple APs occur. While the proportions of spiking patterns in each pool (in parentheses) are relatively similar, it appears that adapting firing patterns are roughly 5 times as common in the H-responder group as S-responders and non-responders. We did not find fastspiking neurons in the H-responder group, though they were found sometimes in S-responders and nonresponders. The PING nomenclature also designates “Intrinsic Burst Spiking” and “Accelerating” spike patterns; however, we did not observe these patterns in the sample (*N* = 55).

| Pattern | H-responders (N = 11)<br>Count (% of column) | S-responders (N = 10)<br>Count (% of column) | Nonresponders (N = 34)<br>Count (% of column) |
| --- | --- | --- | --- |
| Fastspiking | 0 (0%) | 2 (20%) | 8 (23.5%) |
| Nonadapting,<br>nonfastspiking | 3 (27.2%) | 6 (60%) | 16 (47.1%) |
| Adapting | 6 (54.5%) | 1 (10%) | 4 (11.8%) |
| Irregular Spiking | 2 (18.2%) | 1 (10%) | 6 (17.6%) |

Our data also suggest that some of the S or H-responders may have features consistent with VIPergic interneurons of the neocortex based on reductions in inhibitory postsynaptic potentials that we observed during recording. These experiments did not include the use of GABAA and GABAB inhibitors, and IPSPs could therefore persist through the spontaneous activity of the interneurons. Although we did not analyze these events, we noticed reductions in IPSPs occurring after addition of DAMGO in some interneurons, including some H-responders (data not shown, though partially visible off-pulse in Figure 10). The VIPergic interneurons tend to inhibit other interneurons, which may have been manifesting our data (Fu et al., 2014; Jackson, Ayzenshtat, Karnani, & Yuste, 2016; Pi et al., 2013). It is interesting to note that previous research in neocortical slices has found a high degree of overlap between VIP and μOR immunoreactivities (Taki et al., 2000). Electrophysiological studies have also found high rates of VIP expression amongst interneurons that hyperpolarize in response to DAMGO, including neurons that would have been classified here as having Adapting spiking patterns (Férézou et al., 2007). While speculative, this may hint at a relationship between the H and S responders and VIPergic neurons of the neocortex.

**Figure 10.**
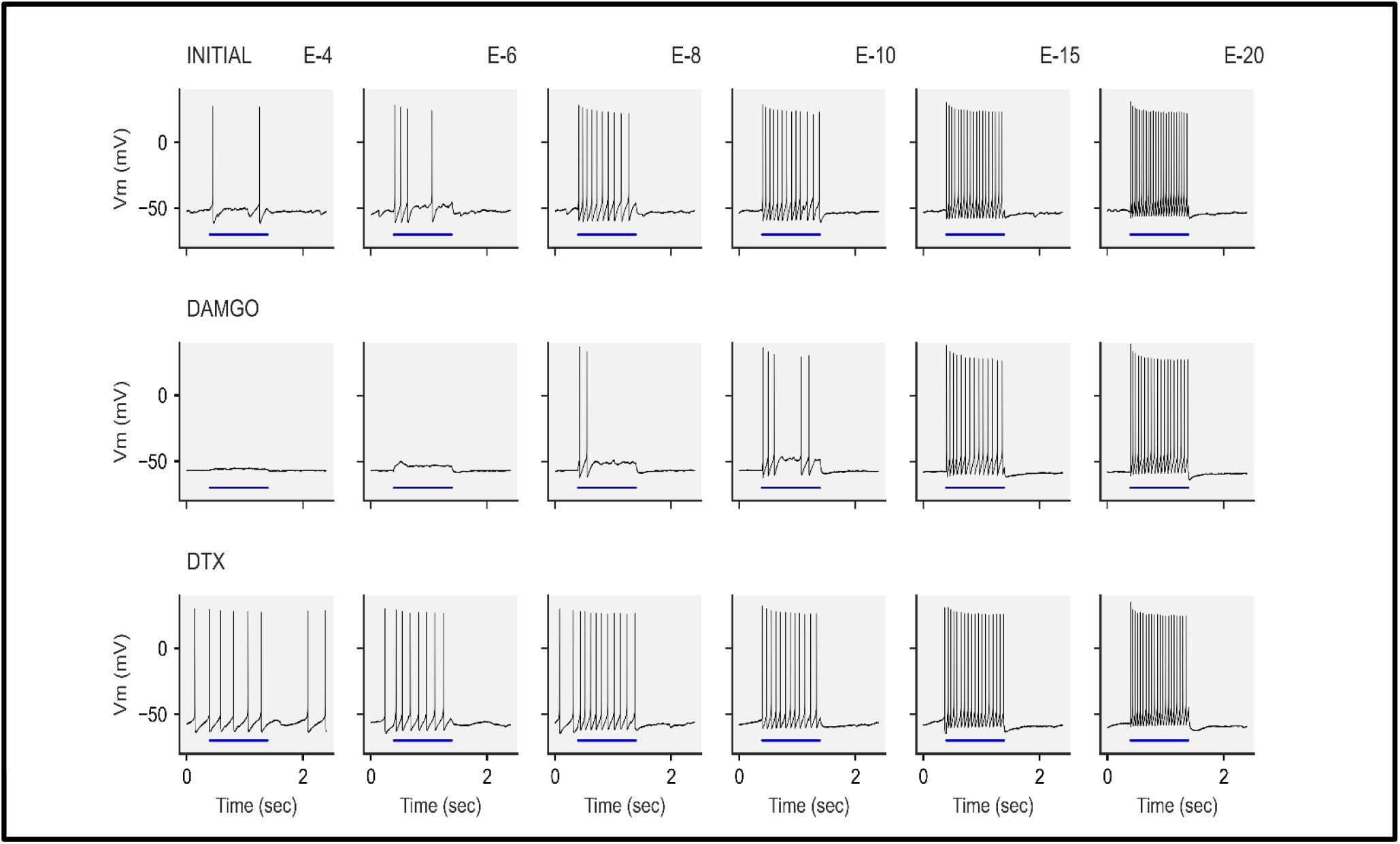
Example of αDTX reversal of DAMGO. A trace representing αDTX reversing effects of DAMGO in AP number and interspike interval. Respectively, each row represents the initial, post-DAMGO, post-αDTX+DAMGO timeslots. Each column represents one of the 20 sequential episodes as the current applications progressively become more depolarizing. For example, E4 is the 4^th^ episode of stimulation, which generally was around AP threshold, while E20 is the final and most-depolarizing of the 20 episodes of the stimulation. The blue horizontal bars below the APs represent the period of 1 second where the depolarizing current was being applied. (Top Row) This neuron began firing APs in the early, less-depolarizing current applications. Application of DAMGO (Middle Row) resulted in the neuron firing APs only at later episodes. (Bottom Row) After addition of αDTX, the neuron once again began firing at earlier episodes and more frequently than it had after exposure to only DAMGO. A slight RMP hyperpolarization after DAMGO is evident here, which was not reversed by αDTX.

**Figure 11.**
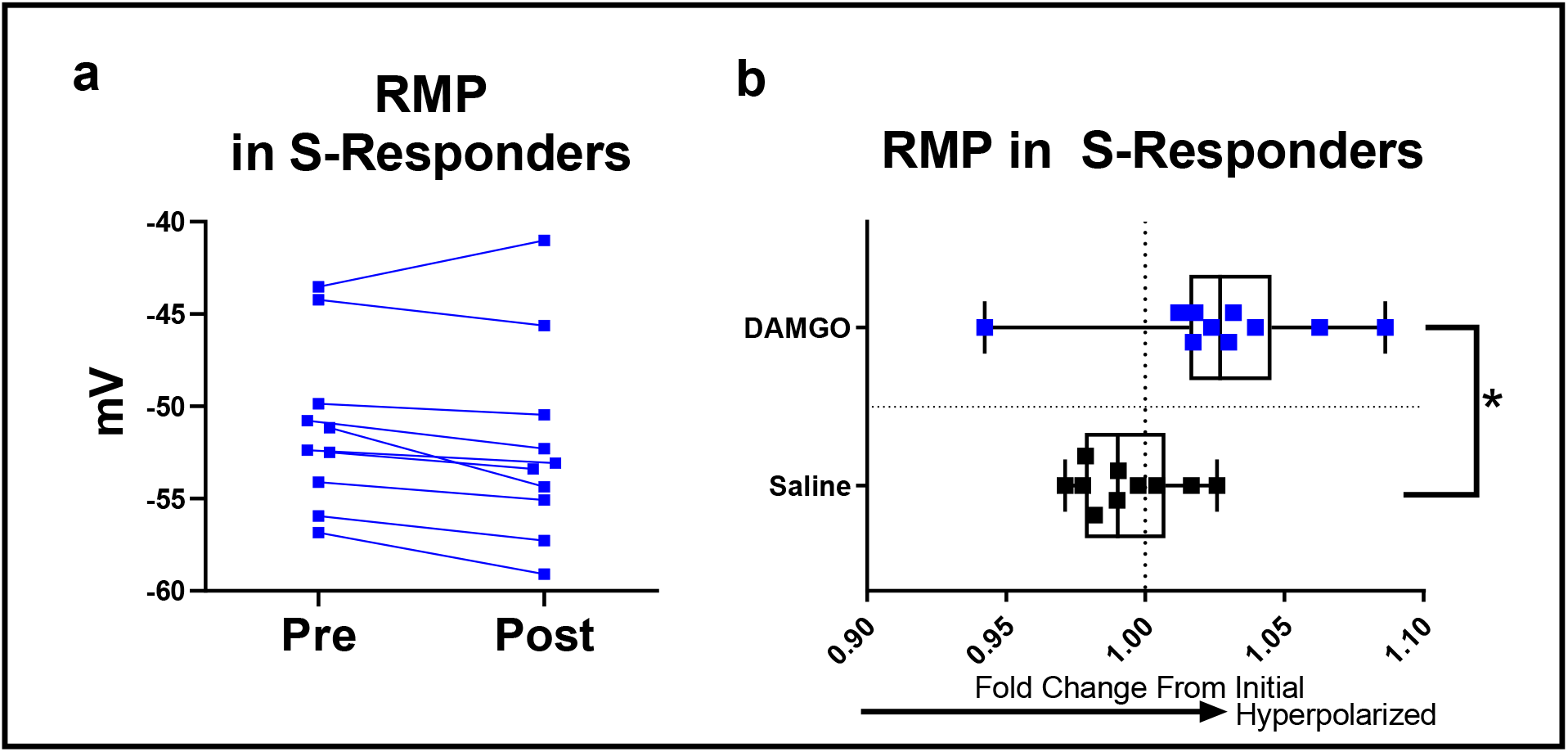
S-responders also hyperpolarize. We compared the fold change in RMP in the (N = 10) S-responders versus random saline to determine if hyperpolarization is found in this group as well. (a) Real values of resting membrane potential in S-responders before and after (3 μM) DAMGO. (b) Fold changes in resting membrane potential in S-responders versus random saline controls (n = 10). We found that the S-responders too underwent a significant hyperpolarization after exposure to DAMGO (*p* < 0.05) compared to the polarization change in saline controls during the same period. Edges of the boxes are drawn at the 1^st^ and 3^rd^ quartiles while the “whiskers” connect the largest and smallest values for that group. We performed statistical testing by confirming distribution was non-significantly different from a normal distribution with the D’Agostino-Pearson omnibus K2 test, and then an unpaired t-test against the fold change of the saline controls.

**Figure 12.**
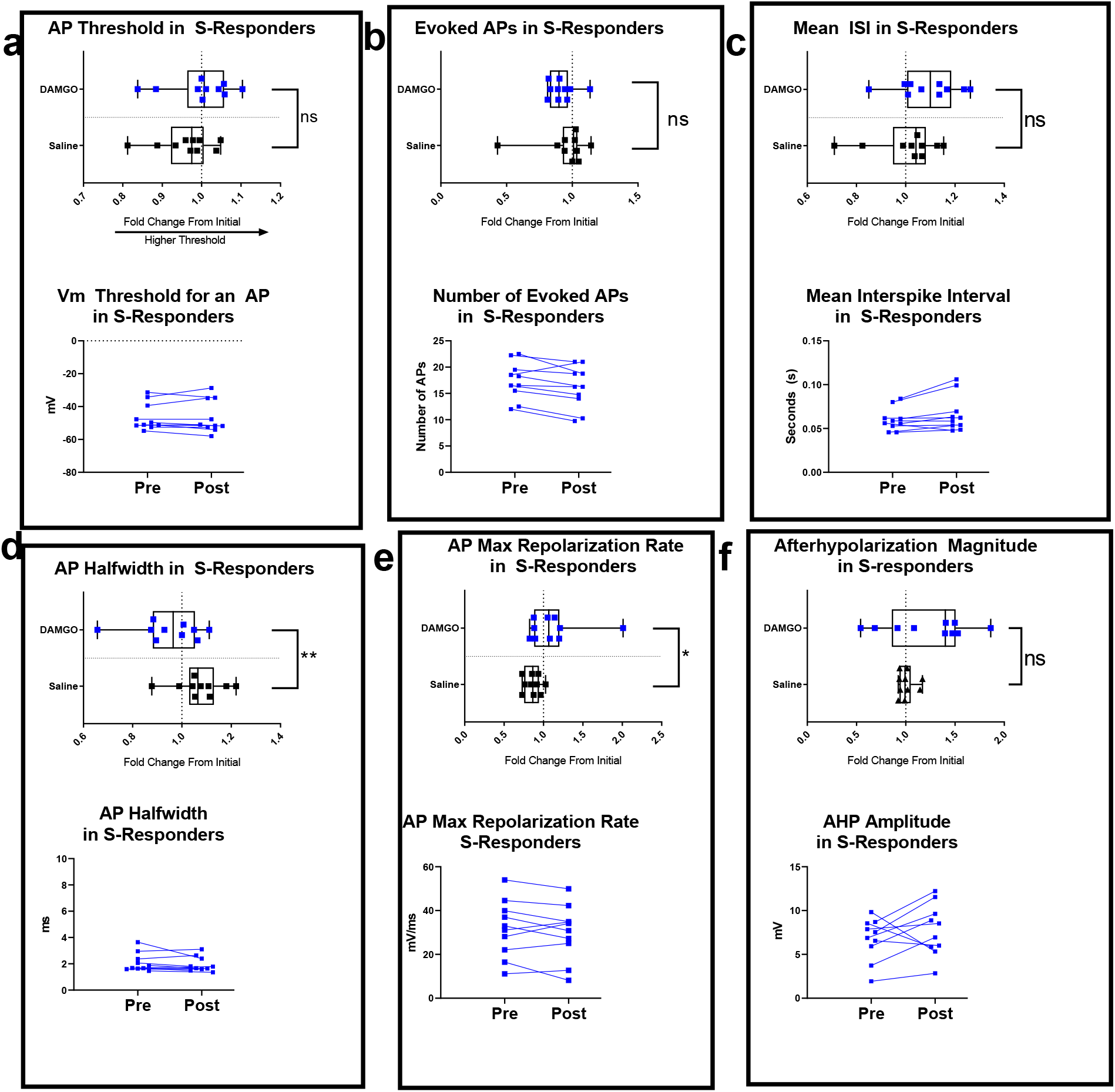
S-responders have changes in AP waveform. We compared the change in each parameter in a sample (*n* = 10) saline group versus the (*N* = 10) S-responders. Boxes are drawn around each set of real values for each measure (bottom of each box), and the fold change (top of each box). We found that these S-responders had significant reductions after DAMGO in (d) halfwidths and significant increases in (e) maximum rates of repolarization compared to saline controls (*p* < 0.05). However, we did not find significant differences in the changes in (a) AP threshold, (b) number of evoked APs, (c) mean interspike interval, or (f) afterhyperpolarization magnitude (*p* > 0.05). Edges are drawn at the 1^st^ and 3^rd^ quartiles while the “whiskers” connect the largest and smallest values for that group. Prior to statistical testing we tested for normality with D’Agostino-Pearson omnibus K2 test (*p* > 0.05). In most instances, we performed statistical testing with unpaired t-tests (one-tailed). However, number of evoked APs and maximum repolarization rates were nonnormally distributed (*p* < 0.05) and we therefore tested those with Mann-Whitney U tests (one-tailed).

It is also important to point out that H and S-responders may not represent discrete populations of interneurons despite the different methodologies to distinguish them from nonresponders; numerous effects we found in H-responders fell just short of significance in the S-responders (Table 6). For example, many S-responders had particularly robust changes in AHP amplitude, which was not captured in the group mean. Although the S-responders generally had weaker responses to DAMGO (Table 7), significant differences in this group may have been more difficult to resolve due to lower samples sizes (N = 10), versus the H-responders (N = 11).

We also found that neurons within the H-responder group possess a variety of spiking patterns (Table 2) and also a range of responses to both DAMGO and αDTX (Figure 9). Within the hyperpolarizing DAMGO responders, we found a spectrum of changes in their AP kinetic parameters; some neurons had pronounced changes in their parameters, and some changed barely at all despite their greater hyperpolarization (Figures 9 & 14). The assortment of ion channels in each particular neuron may have led to varied responses to DAMGO, but variability may also have resulted from the complex intracellular signaling cascades of GPCRs; μORs are shown in non-cortical regions to modulate αDTX-sensitive currents through the Gαi subunit, phospholipase A2 and the lipoxygenase pathway (Faber & Sah, 2004; Vaughan et al., 1997), meanwhile upregulation of GIRK channels appears to be mediated through direct interaction by activated Gβγ subunits (Huang, Slesinger, Casey, Jan, & Jan, 1995; Inanobe et al., 1995; Logothetis, Kurachi, Galper, Neer, & Clapham, 1987; Wickman et al., 1994). It is therefore conceivable that neurons have a variety of responses to DAMGO ranging from activating GIRK channels, to activating voltage-sensitive currents, based on diverging signaling cascades.

The neurons in the H and S-responder categories were identified using two different measures (RMP and spontaneous APs, respectively), which themselves can be manifestations of different target ion channels of μORs, or different localization of the μORs among the neuron types. Studies in the cortex suggest that hyperpolarization is not always found in a DAMGO response, and that there is incomplete overlap between GIRK channels and μORs (Bausch et al., 1995; Wimpey & Chavkin, 1991). Therefore, larger hyperpolarization may not always be seen in μOR+ neurons if GIRK channels are not involved. However, the drop in spontaneous APs seen in the S-responders is another interesting feature. Studies of this receptor in neocortical interneurons using DAMGO have found reductions in non-elicited APs, which appear to originate in distal processes of some interneurons (Krook-Magnuson et al., 2011; Suzuki et al., 2014). These mechanisms illustrate that the μOR has several known mechanisms to downregulate interneurons which can vary among the interneuronal types.

We found a range of AP kinetics as well as firing frequency properties (Table 7) are altered by the μOR. We found that the AP kinetics were not likely to be mediated by the αDTX-sensitive channels, though measures of AP probability (ISI, AP number, and AP threshold) were related to the αDTX-sensitive current (Figure 8 and Table 5). We found that αDTX frequently reversed (Figure 9) DAMGO-induced changes in spiking frequency in the H-responders with a wide spectrum of responses, with neurons that underwent DAMGO-induced changes in ISI and evoked APs usually being partially or fully reversed to baseline. Interestingly αDTX did seem to have an effect in 2 or 3 neurons in RMP and AP halfwidth which was not captured by the mean, however it is difficult to say whether this was a genuine reaction to the drug by certain subclasses of interneurons, or if they were simply outliers in the data (Figure 9).

It is interesting that αDTX reversed DAMGO effects in ISI and number of evoked APs without reversing any of the AP kinetic features that we measured. The mechanism behind the αDTX-sensitive current suppression of follow-up APs is not immediately clear considering that changes in AHP magnitude was not significantly changed by DAMGO nor αDTX. In other parts of the brain, this current is responsible for modulating spike frequency adaptation – a feature that limits follow-up APs during prolonged depolarizations (Faber & Sah, 2004). It appears therefore possible that, based on DAMGO’s effect and αDTX’s (usually) partial reversal of it (Figure 9) that other voltage-sensitive potassium currents are being modulated by μORs. For example, the Kv3 and Kv4 family potassium channels are found in cortical neurons and are implicated in modulating these properties (Burkhalter, Gonchar, Mellor, & Nerbonne, 2006; Carrasquillo, Burkhalter, & Nerbonne, 2012; Martina, Schultz, Ehmke, Monyer, & Jonas, 1998; Rudy et al., 2009). The relationship of AP threshold with the μOR and αDTX-sensitive current was a particularly nuanced feature in hyperpolarizing DAMGO responders and appears to track with prior findings from the BLA where μORs modulate αDTX-sensitive current without changing the V_m_ threshold for an AP (Faber & Sah, 2004). Here, αDTX produced a hyperpolarizing shift in AP threshold despite the μOR’s apparent noninfluence over AP threshold (Figures 6c and 9b). Yet the μOR still appears to regulate αDTX current; μOR stimulation with DAMGO suppressed follow-up APs, which was partially reversed by αDTX. At face value, it appears that μOR+ interneurons have αDTX-sensitive currents at rest that influence AP threshold, yet these currents can be upregulated to suppress APs after μOR-stimulation. One explanation is that αDTX-sensitive channels are expressed in areas that the μOR is not localized to. For instance, αDTX-sensitive channels in axon initial segments of cortical neurons can modulate AP threshold (Inda, DeFelipe, & Muñoz, 2006; Kole, Letzkus, & Stuart, 2007), but if the μORs localize to soma and dendrites, they may very well modulate somatic channels, but not AIS-localized ion channels, where they may be already functioning without DAMGO stimulation.

One drawback of this study is that we did not test the inverse drug exposure of αDTX before DAMGO and therefore it is difficult to assess the baseline function of the αDTX-sensitive channels in DAMGO-responders to directly relate μOR-stimulation with the upregulation of αDTX-sensitive channels. When designing these experiments, we believed it might have been possible that αDTX would block DAMGO-induced hyperpolarization (the primary indicator of a positive DAMGO response) and therefore prevent us from ultimately identifying H-responders. However, we found that αDTX usually failed to reverse DAMGO effects, including hyperpolarization. The inverse drug sequence may be the most ideal way to measure baseline αDTX activity in DAMGO responders. On the other hand, there were some neurons that did undergo noticeable depolarization and had pronounced increases in spontaneous APs, and we may have overlooked several H-responders and S-responders had we used inverse drug exposures (Figure 9a).

It may be argued that the hyperpolarizing effect of DAMGO found in the hyperpolarizing responders and, to a lesser degree, in spontaneous DAMGO responders may have influenced action potential parameters. This is unlikely for several reasons. First, most of the action potential analyses were done at several current steps above threshold and we used the average values for APs 3-5 in those trains, thereby mitigating the influence of hyperpolarized RMP on action potential parameters. In addition, most of the APs we observed were initiated from approximately the same V_m_ regardless of their change in RMP (for example, Figure 10). Second, αDTX reversed DAMGO-induced changes in ISI and number of evoked APs in hyperpolarizing DAMGO responders despite αDTX’s failure to reverse DAMGO-induced hyperpolarization (Table 3 and Figure 8a) and broad range of other features in the post-hoc analyses (Table 9). Third, we compared DAMGO depolarizers (predicted to have had no μOR response) against neurons that had hyperpolarized slightly in the saline group; this comparison “flipped the sign” to the saline group being significantly more hyperpolarized then the DAMGO group. However, in all measures resulting from this comparison, we found no significant differences in action potential parameters, ISI, and number of evoked action potentials (Figure 13). This analysis suggests that the parameters we were investigating during the current steps were not being influenced by the neuron’s RMP regardless if the neuron had depolarized under the DAMGO challenge.

**Figure 13.**
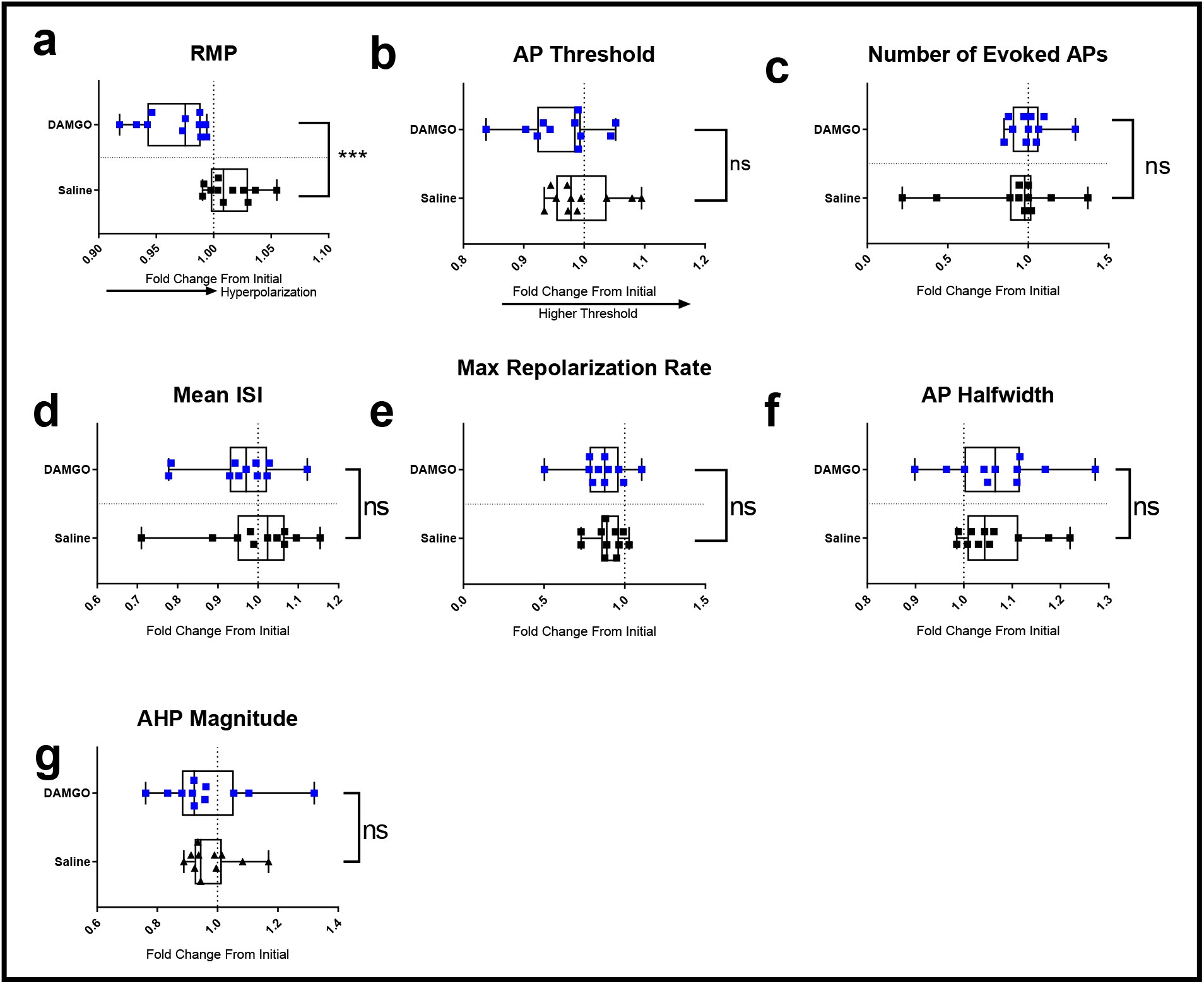
Polarity shift does not account for other changes. In these analyses, we compared neurons that depolarized in the (*n* = 11) DAMGO condition and neurons that hyperpolarized in the (*n* = 11) saline control. These changes were expected to be due to stochastic fluctuations in their RMP during this period, and we therefore expected null results. We selected these neurons based on their changes in (a) RMP which, predictably, had significantly depolarized in the DAMGO condition relative to the change in saline controls (*p* < 0.05). However, none of the other changes we tested were significantly different from controls (*p* > 0.05); we did not find significant differences in their changes of (b) AP threshold, (c) number of evoked APs, (d) interspike interval, (e) maximum repolarization rates, (f) AP halfwidth, nor (g) magnitude of their hyperpolarization when the groups were compared to controls in the same manner as the H and S-responders were. Comparisons of the DAMGO-depolarizers versus saline-hyperpolarizers were made with unpaired t-tests (one-tailed) and summarized on Table 8. Edges of the boxes are drawn at the 1^st^ and 3^rd^ quartiles while the “whiskers” connect the largest and smallest values for that group. We performed statistical testing with unpaired t-tests (one-tailed) after testing for skewness and kurtosis with D’Agostino-Pearson omnibus K2 test (*p* > 0.05). Only halfwidth and maximum repolarization rates were significantly different (*p* < 0.05) from a normal distribution, and therefore tested with Mann-Whitney U tests.

**Figure 14.**
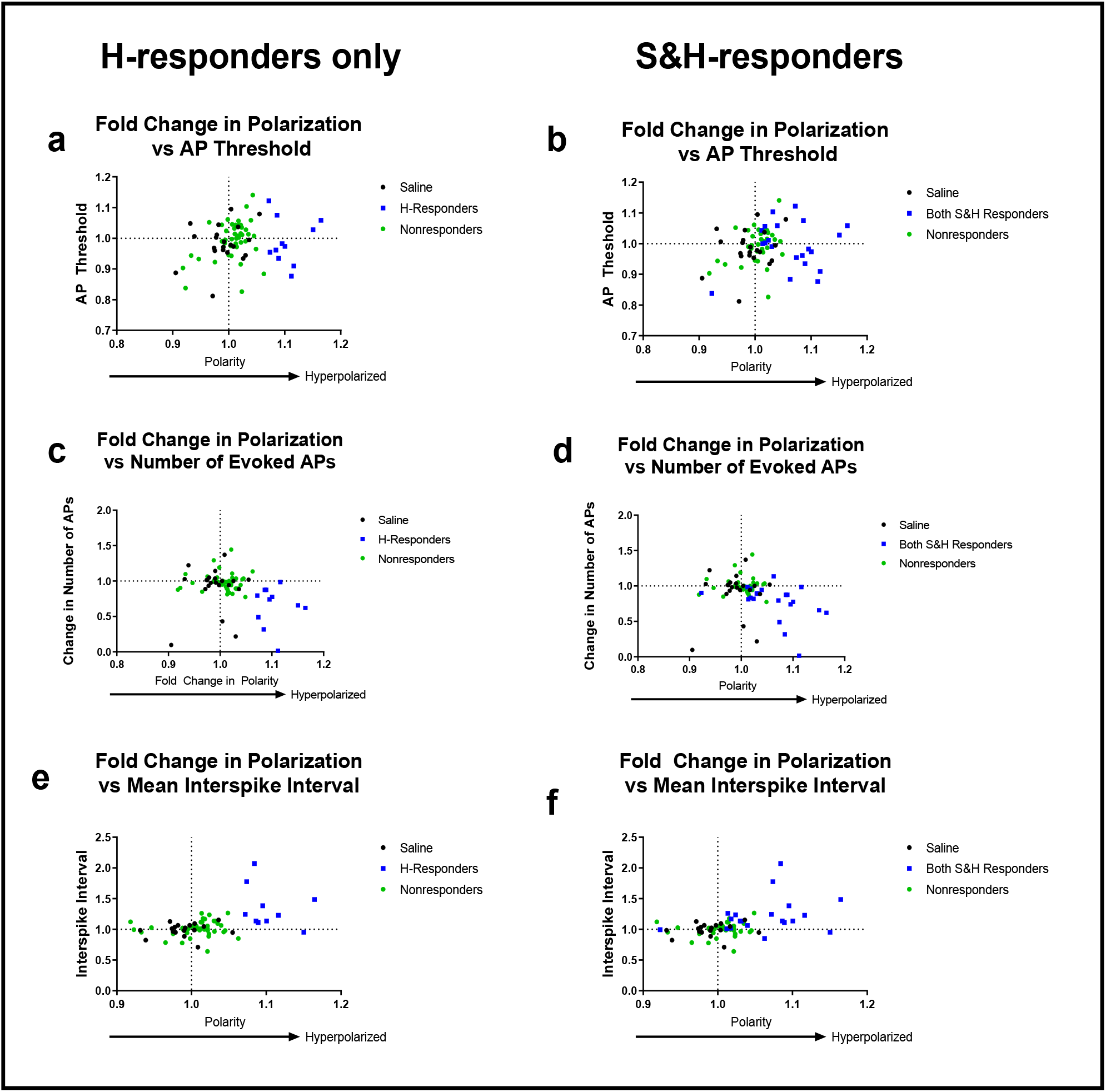
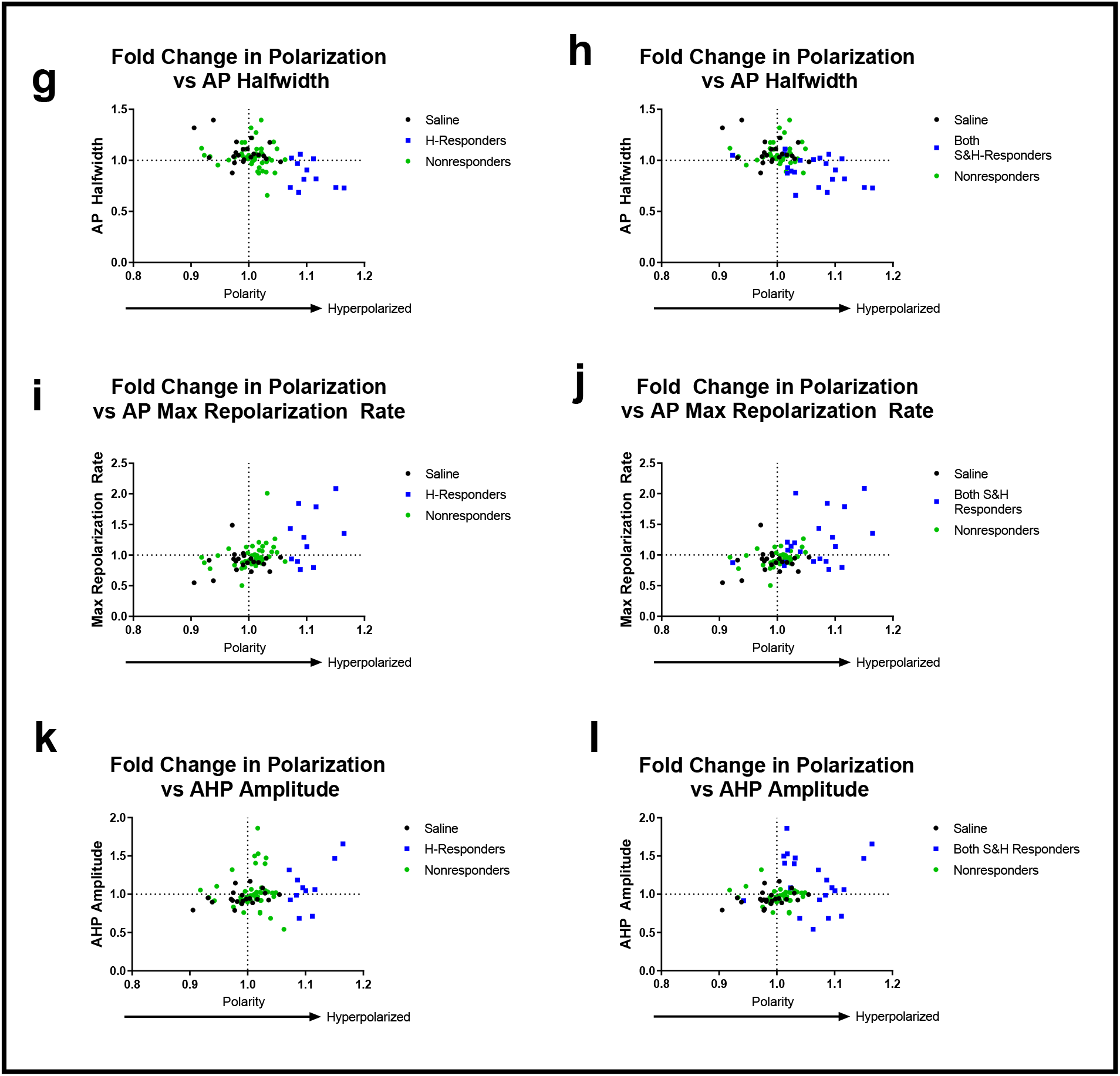
Responder categories captured most large changes. To determine whether the changes we were attempting to detect had successfully been predicted by the S and H responder groups, we compared polarization (x ais) with the changes in AP parameters we were measuring (y axis).Vertical displacement on the y axis therefore indicates that the neuron had large changes in the measure being compared and can be compared with its polarization (horizontal displacement). Left column only includes the H-responders (blue squares), while the right column expands the responder category to both S&H-responders (blue squares). Saline-controls (black) are included in these images to illustrate the pattern of stochastic or entropic changes that should be considered in interpretations of these graphs. (a,b) AP Threshold versus polarization. We previously found no DAMGO effect in this measure, and the distribution shows scattering around the origin. (c,d) Number of evoked APs show a downward and right skew, as we expected the DAMGO responders were firing fewer evoked APs. (e,f) Interspike interval showed and upwards and right skew in the responders, as we expected, since a DAMGO response should reduce spike frequency (and increase temporal spacing). (g,h) AP halfwidth showed a downwards and right skew, particularly in DAMGO responders. (i,j) AP maximum repolarization rates show and upwards and right skew in the DAMGO responders. (k,l) Afterhyperpolarization amplitude versus polarization shows an upward spread of S and H-responders.

**Figure 15.**
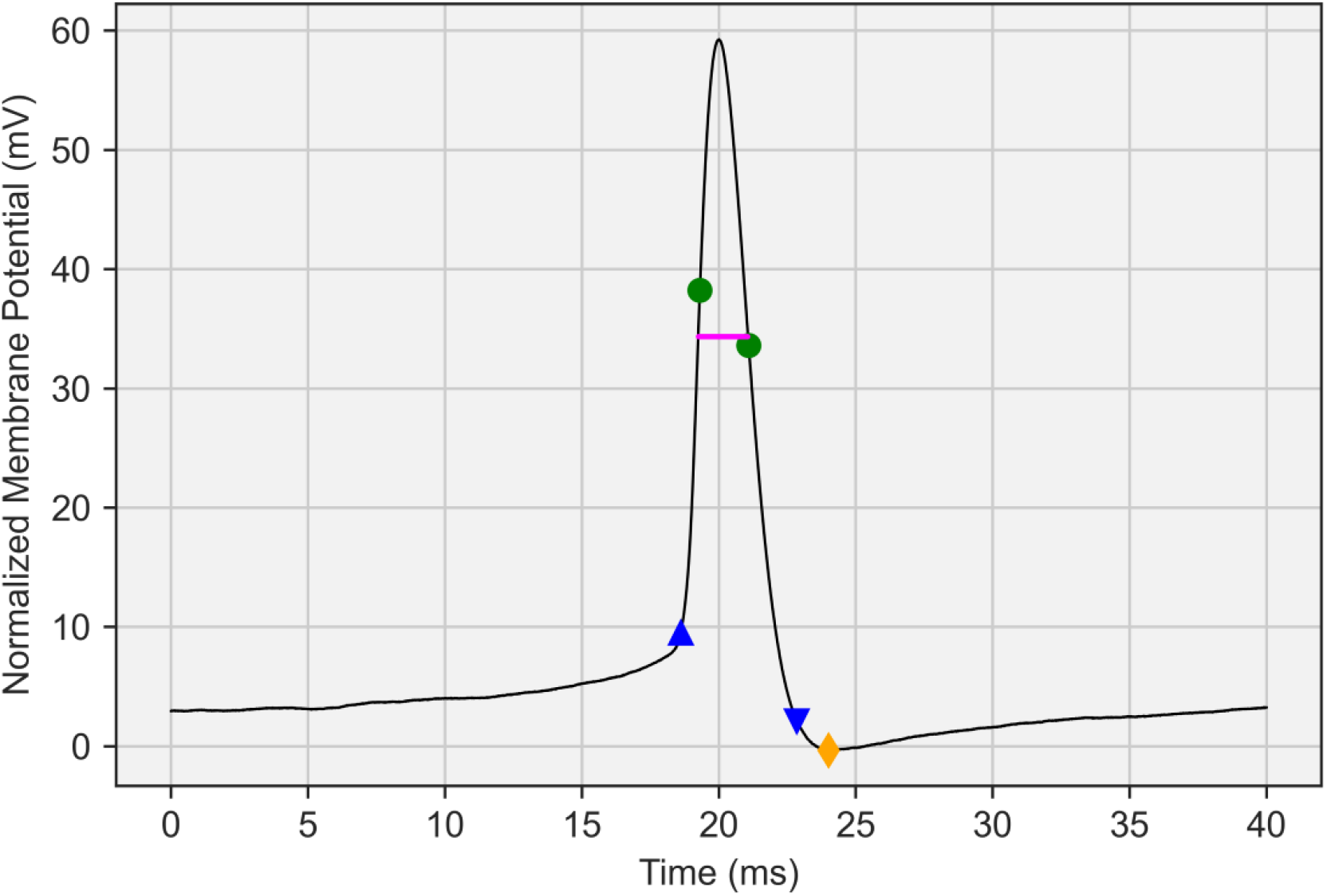
Sample AP diagram. AP kinetics were measured by the script and an example is shown here to illustrate the derivation of the measurements (Upward-facing blue arrow) Start of the AP demarcated at 10% of the maximum depolarization rate. V_m_ at this point represents AP threshold. (First Green Circle) Maximum rate of depolarization of the AP. (Second Green Circle) Maximum rate of repolarization of the AP. (Downward-facing blue arrow) 10% of maximum rate of repolarization, marking the end of the AP. (Yellow Diamond) Lowest point of the afterhyperpolarization. Afterhyperpolarizations were measured as the difference between the y value of the start of the AP with the lowest point of the afterhyperpolarization, out to a maximum distance of 2ms from the point where the repolarization phase returns to the same V_m_ as the start of the AP at the upward blue arrow. (Pink Horizontal Line) The halfwidth of the AP was measured at half the height of the AP from the threshold V_m_ of the AP at the upward blue arrow.

**Table 3.**
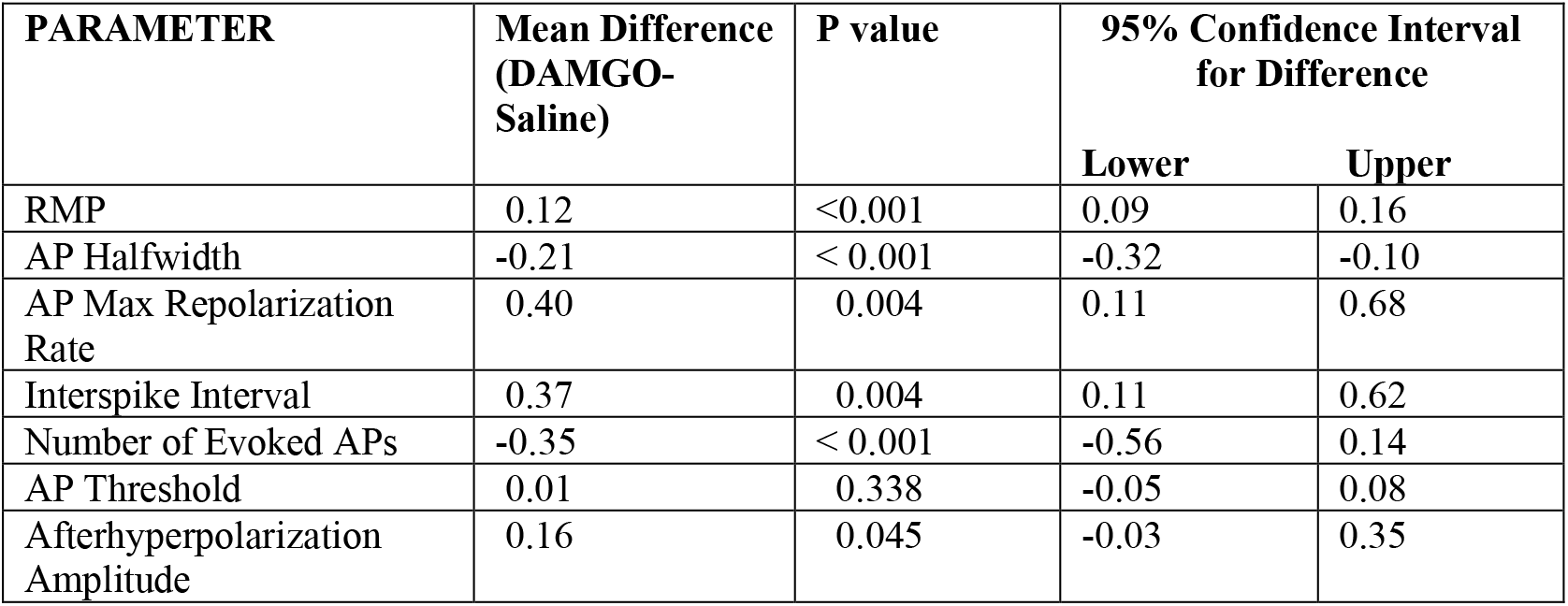
Summary of DAMGO effects in H-responders. We compiled a statistical summary and effect sizes for H-responders in all 7 measures that we were primarily investigating. The change in each property was derived by dividing post-DAMGO/pre-DAMGO values (i.e., slot 2/slot 1) to calculate the fold change. Here, these fold changes in the saline group (expected to represent the change due to time) was subtracted from the fold change in the DAMGO group to derive the effect of DAMGO. Each mean difference is a proportion of one (e.g., a mean difference of −0.21 is a reduction of 21% in the DAMGO condition over the saline control). Comparisons of the change in saline and DAMGO group were tested first for normality with the D’Agostino-Pearson omnibus K2 test. Normally-distributed data were tested for significance (α = 0.05) with one-tailed unpaired t-tests and nonnormally-distributed data were tested with one-tailed Mann-Whitney U tests.

| PARAMETER | Mean Difference<br>(DAMGO-Saline) | P value | 95% Confidence Interval<br>for Difference |  |
| --- | --- | --- | --- | --- |
|  |  |  | Lower | Upper |
| RMP | 0.12 | <0.001 | 0.09 | 0.16 |
| AP Halfwidth | -0.21 | < 0.001 | -0.32 | -0.10 |
| AP Max Repolarization Rate | 0.40 | 0.004 | 0.11 | 0.68 |
| Interspike Interval | 0.37 | 0.004 | 0.11 | 0.62 |
| Number of Evoked APs | -0.35 | < 0.001 | -0.56 | 0.14 |
| AP Threshold | 0.01 | 0.338 | -0.05 | 0.08 |
| Afterhyperpolarization Amplitude | 0.16 | 0.045 | -0.03 | 0.35 |

**Table 4.** Few H-responders shift spiking patterns after DAMGO. We identified the spiking patterns of H-responders based on PING categories before and after DAMGO. Neurons are ordered by row arbitrarily. Boldtype rows are neurons that change firing type. The majority of the neurons remained the same spiking patterns, though 3 had changed after DAMGO.

| Pre-DAMGO Pattern | Post-DAMGO Pattern |
| --- | --- |
| Adapting | Adapting |
| Adapting | Adapting |
| Adapting | Adapting |
| Adapting | Adapting |
| Adapting | Adapting |
| **Adapting** | **Irregular Spiking** |
| Irregular Spiking | Irregular Spiking |
| **Irregular Spiking** | **Adapting** |
| Non-adapting, Non-fastspiking | Non-adapting, Non-fastspiking |
| Non-adapting, Non-fastspiking | Non-adapting, Non-fastspiking |
| **Non-adapting, Non-fastspiking** | **Adapting** |

**Table 5.**
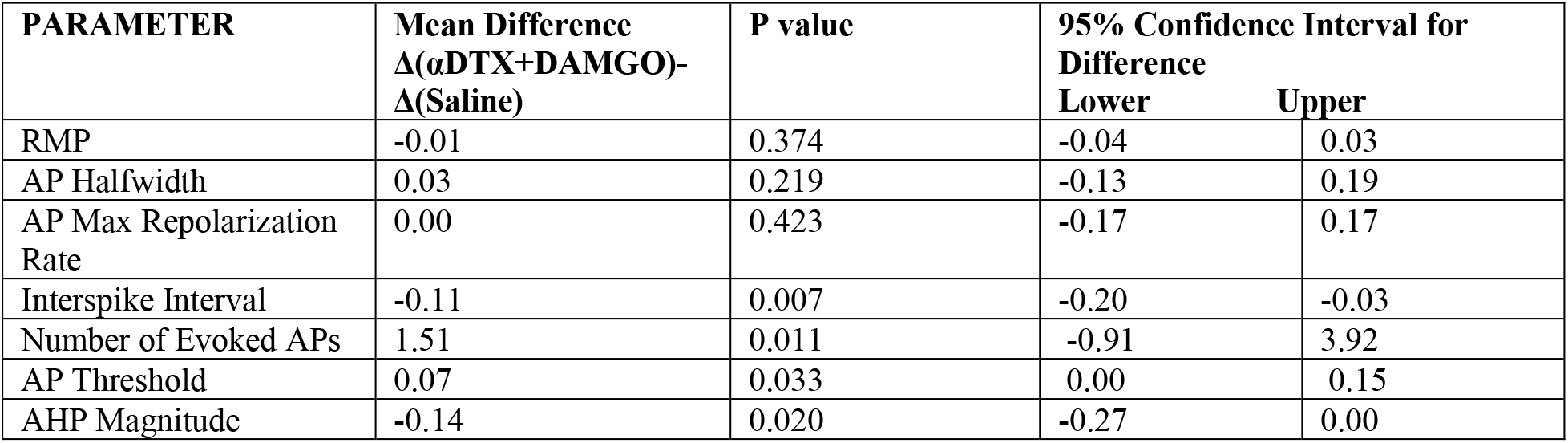
αDTX-sensitive channels reverse some DAMGO effects. Mean values in this summary were derived by subtracting the mean fold change in the saline group from the mean fold change in the DAMGO group. The fold changes for each of the groups were compared with unpaired t tests (one-tailed) for significance or Mann-Whitney U-tests.

| PARAMETER | Mean Difference<br>$\Delta(\alpha\text{DTX}+\text{DAMGO})-$<br>$\Delta(\text{Saline})$ | P value | 95% Confidence Interval for Difference | |
| --- | --- | --- | --- | --- |
|  |  |  | Lower | Upper |
| RMP | -0.01 | 0.374 | -0.04 | 0.03 |
| AP Halfwidth | 0.03 | 0.219 | -0.13 | 0.19 |
| AP Max Repolarization Rate | 0.00 | 0.423 | -0.17 | 0.17 |
| Interspike Interval | -0.11 | 0.007 | -0.20 | -0.03 |
| Number of Evoked APs | 1.51 | 0.011 | -0.91 | 3.92 |
| AP Threshold | 0.07 | 0.033 | 0.00 | 0.15 |
| AHP Magnitude | -0.14 | 0.020 | -0.27 | 0.00 |

**Table 6.** DAMGO Effect in S-responders. A statistical summary and estimates of effect sizes for the spontaneous-AP responders (S-responders). The change in each property was derived by dividing the post-DAMGO/pre-DAMGO values (i.e., slot 2/slot 1) to calculate the proportion of the change. For instance, a change of 0.28 in AP maximum repolarization rate is an average increase of 28% in the DAMGO S-responders. Comparisons of the change in saline and DAMGO group were performed by unpaired t-tests (one tailed). Values for saline were subtracted from DAMGO values to calculate the true effect of the drug (and subtract the effect of time). Prior to statistical testing, we tested for normality with D’Agostino-Pearson omnibus K2 test. In most instances, we performed statistical testing with unpaired t-tests (one-tailed) (*p* > 0.05). However, number of evoked APs and maximum repolarization rates were nonnormally distributed (*p* < 0.05) and we therefore tested those with Mann-Whitney U tests (one-tailed) instead.

| PARAMETER | Mean Difference<br>(DAMGO-Saline) | P value | 95% Confidence Interval for<br>Difference |  |
| --- | --- | --- | --- | --- |
|  |  |  | Lower | Upper |
| RMP | 0.03 | 0.003 | 0.00 | 0.06 |
| AP Halfwidth | -0.12 | 0.013 | -0.23 | -0.02 |
| AP Max Repolarization<br>Rate | 0.25 | 0.009 | 0.02 | 0.49 |
| Interspike Interval | 0.08 | 0.086 | -0.04 | 0.21 |
| Number of Evoked APs | -0.03 | 0.072 | -0.03 | 0.07 |
| AP Threshold | 0.04 | 0.141 | -0.03 | 0.10 |
| Afterhyperpolarization<br>Amplitude | 0.23 | 0.051 | -0.05 | 0.52 |

**Table 7.** Summary of effects for H-responders vs S-responders. To provide a side-by-side comparison of H and S-responders and compare their response to DAMGO, we complied data from H-responders (Table 2) and data from S-responders (Table 3) and re-display them in this table. Numerical values reflect the mean fold change in that particular responder group (see top row) for each parameter (left column). Mean differences were derived by subtracting fold change in the saline group from the DAMGO responder group to isolate the effect of DAMGO from the effect of time. For example, a value of 0.40 for H-responder AP max repolarization rate reflects a 40% increase for maximum rate of AP repolarization in H-responders compared to the control, which trended larger than the mean 28% change in the S-responders. Only significant (p < 0.05) values are shown in the table; cells are intentionally left blank when effects were nonsignificant (Unpaired t-test or Mann-Whitney U test, as described in Tables 3 and 6). We found that the H-responders had a wider range of detected effects to DAMGO, as well as having larger effect sizes.

| PARAMETER | Mean Difference H-responder<br>(DAMGO-Saline) | Mean Difference S-responder<br>(DAMGO-Saline) |
| --- | --- | --- |
| RMP | 0.12 | 0.03 |
| AP Halfwidth | -0.21 | -0.12 |
| AP Max Repolarization Rate | 0.40 | 0.25 |
| Interspike Interval | 0.37 | - |
| Number of Evoked APs | -0.35 | - |
| AP Threshold | - | - |
| Afterhyperpolarization Amplitude | 0.16 | - |

**Table 8.** Comparisons for the effect of polarity shift. Neurons that depolarized in the DAMGO condition were compared against the most hyperpolarizing neurons in the saline condition. This table compares the mean changes in each measure for these groups. Statistical testing revealed no significant changes in any parameter we compared, other than the RMP which was the basis for their selection to these groups. Change in RMP was significantly different from the controls (*t*(20) = 4.61, *p* < 0.001), however changes in AP halfwidth (*t*(20) = 0.24, *p* = 0.405), max repolarization rate (*t*(20) =0.25, *p* = 0.245), interspike interval (*U* = 43, *p* = 0.135), number of evoked APs (*t*(20) = 1.05, *p* = 0.154), AP threshold (*t*(20) = 0.70, *p* = 0.245), and AHP (*U* = 48, *p* = 0.219) were nonsignificantly (*p* < 0.05) different from each other. Prior to statistical comparisons, distributions were tested with D’Agostino-Pearson omnibus K2 test. Normally-distributed data (*p* > 0.05) were tested with unpaired t-tests (one-tailed) while nonnormally-distributed data were tested with Mann-Whitney U tests (one-tailed).

| PARAMETER | Mean Difference<br>(DAMGO-Saline) | P value | 95% Confidence Interval for<br>Difference |  |
| --- | --- | --- | --- | --- |
|  |  |  | Lower | Upper |
| RMP | -0.05 | <0.001 | -0.07 | 0.03 |
| AP Halfwidth | -0.02 | 0.405 | -0.08 | 0.09 |
| AP Max Repolarization<br>Rate | -0.04 | 0.245 | -0.15 | 0.08 |
| Interspike Interval | -0.04 | 0.135 | -0.04 | 0.05 |
| Number of Evoked APs | 0.11 | 0.154 | -0.11 | 0.32 |
| AP Threshold | -0.03 | 0.104 | -0.08 | 0.02 |
| AHP Amplitude | -0.01 | 0.219 | -0.12 | 0.09 |

**Table 9.** Our Python script analyzed 44 measures and 5 ratios, and only 7 have been discussed previously here. We therefore analyzed the 44 remaining measures and 5 ratios not presented yet to determine whether DAMGO or αDTX were have significant effects on parameters that have not yet been discussed here. Several mixed-model repeated measures MANOVAs were repeated for every subcategory (left column) for both DAMGO and αDTX to identify Slot*Drug interaction effects which may indicate that those drugs were modulating the properties listed on the left-side column H-responders (*N* = 11) were compared with saline controls (*N* = 21) on various parameters for a Slot*Group interaction that would suggest a drug (DAMGO or αDTX) that was independent of degradation reflected in the saline control group. Timeslots analyzed were Initial → DAMGO for DAMGO effects, and DAMGO → αDTX+DAMGO for αDTX effects in a background of DAMGO. In order to determine whether these drugs acted only specifically on certain APs (e.gs., only the first AP, second AP, or only APs after that), we analyzed these APs separately for Slot*Group interactions. Boldtype are significant effects. In membrane and threshold properties, omnibus tests revealed a significant Time*Group interaction for DAMGO (*F*(3,26) = 11.42, *p* < 0.001, *Wilks’ λ* = 0.568) in the 3 combined measures, however no significant αDTX effects were found in the combined variables (*F*(3,25) = 0.36, *p* = 0.784, *Wilks’ λ* = 0.041). An omnibus test on parameters of the first AP properties found a significant Slot*Group interaction for DAMGO effects on the first AP in trains (*F*(11,20) = 11.31, *p* <0.001, *Wilks’ λ* = 0.861), but not the omnibus test for changes in AP1 of the αDTX timeslots (*F*(11,19) = 1.94, *p* =0.082, *Wilks’ λ* =0.543). The omnibus test for changes in AP2 properties during DAMGO exposure found a significant Slot*Group interaction (*F*(11,19) = 2.53, *p* = 0.036, *Wilks’ λ* = 0.595), but this interaction was not found during αDTX exposure for AP2 (*F*(11,18) = 1.67, *p* = 0.081, *Wilks’ λ* = 0.559). Finally, we tested for drug effects by averaging all APs after the second AP (AP3+) and comparing that change against the change in the saline control, and we found a significant Slot*Group interaction in those properties for DAMGO (*F*(9,22) = 6.31, *p* = 0.001, *Wilks’ λ* = 0.721) and also for αDTX (*F*(9,21) = 3.16, *p* = 0.014, *Wilks’ λ* = 0.576). We tested for DAMGO effects in the combined variables of interspike properties and found a significant effects (*F*(9,21) = 3.13, *p* = 0.19, *Wilks’ λ =* 0.610), and we also found an effect of αDTX in the variables as well (*F*(9,17) = 2.64, p = 0.040, *Wilks’ λ* = 0.583). Finally, tested the ratios of some of these measures to determine whether some of these measures were changing differently over the course of their series of APs. We found a nonsignificant effect of DAMGO (*F*(5,22) = 1.90, *p* = 0.441, *Wilks’ λ* = 0.185) in these measures as well as a nonsignificant effect for αDTX (*F*(5,21) = 1.20, *p* = 0.342, *Wilks’ λ* = 0.223).

| | PARAMETER | DAMGO<br>p value | DAMGO<br>Full MANOVA<br>Results | αDTX<br>p value | αDTX<br>Full MANOVA<br>Results |
| --- | --- | --- | --- | --- | --- |
| Membrane and $V_m$<br>Threshold Parameters | Input Resistance | 0.064 | $F(1,28) = 3.71$,<br>$\eta_p^2 = 0.117$ | 0.512 | $F(1,27) = 0.44$,<br>$\eta_p^2 = 0.016$ |
| | AP Number at Threshold | 0.321 | $F(1,28) = 1.02$,<br>$\eta_p^2 = 0.035$ | 0.581 | $F(1,27) = 0.31$,<br>$\eta_p^2 = 0.011$ |
| | AP Halfwidth at<br>Threshold | **<0.001** | $F(1,28) = 23.13$,<br>$\eta_p^2 = 0.452$ | 0.536 | $F(1,27) = 0.39$,<br>$\eta_p^2 = 0.014$ |
| First AP Parameters | Maximum Depolarization<br>Rate | **<0.001** | $F(1,30) = 22.47$,<br>$\eta_p^2 = 0.428$ | 0.917 | $F(1,29) = 0.78$,<br>$\eta_p^2 = 0.011$ |
| | Average Depolarization<br>Rate | **0.001** | $F(1,30) = 13.33$,<br>$\eta_p^2 = 0.308$ | 0.472 | $F(1,27) = 0.53$,<br>$\eta_p^2 = 0.018$ |
| | Time to Max<br>Depolarization Rate | **0.001** | $F(1,30) = 12.32$,<br>$\eta_p^2 = 0.291$ | 0.612 | $F(1,27) = 0.26$,<br>$\eta_p^2 = 0.009$ |
| | Time to peak | **0.003** | $F(1,30) = 10.60$,<br>$\eta_p^2 = 0.251$ | 0.785 | $F(1,27) = 0.08$,<br>$\eta_p^2 = 0.003$ |
| | Peak Amplitude | 0.084 | $F(1,30) = 3.19$,<br>$\eta_p^2 = 0.096$ | 0.172 | $F(1,27) = 1.96$,<br>$\eta_p^2 = 0.063$ |
| | Maximum Repolarization<br>Rate | **0.001** | $F(1,30) = 14.32$,<br>$\eta_p^2 = 0.323$ | 0.917 | $F(1,27) = 0.01$,<br>$\eta_p^2 < 0.001$ |
| | Average Repolarization<br>Rate | **0.001** | $F(1,30) = 13.65$,<br>$\eta_p^2 = 0.313$ | 0.716 | $F(1,27) = 0.14$,<br>$\eta_p^2 = 0.005$ |
| | Time to max<br>Repolarization Rate | **0.014** | $F(1,30) = 6.75$,<br>$\eta_p^2 = 0.184$ | 0.949 | $F(1,27) < 0.01$, $\eta_p^2$<br>$< 0.001$ |
| | AP1 Halfwidth | **<0.001** | $F(1,30) = 21.79$,<br>$\eta_p^2 = 0.421$ | 0.341 | $F(1,27) = 0.94$, $\eta_p^2$<br>$= 0.031$ |
| | AP1 90% Width | **<0.001** | $F(1,30) = 21.79$,<br>$\eta_p^2 = 0.421$ | 0.949 | $F(1,27) < 0.01$,<br>$\eta_p^2 < 0.001$ |
| | Afterhyperpolarization<br>Amplitude | **0.042** | $F(1,30) = 4.53$,<br>$\eta_p^2 = 0.131$ | **0.043** | $F(1,27) = 4.48$,<br>$\eta_p^2 = 0.134$ |
| Second AP<br>Parameters | Maximum Depolarization<br>Rate | **0.001** | $F(1,29) = 14.02$,<br>$\eta_p^2 = 0.326$ | 0.462 | $F(1,28) = 0.56$,<br>$\eta_p^2 = 0.019$ |
| | Average Depolarization<br>Rate | **0.001** | $F(1,29) = 15.56$,<br>$\eta_p^2 = 0.334$ | 0.501 | $F(1,28) = 0.47$,<br>$\eta_p^2 = 0.016$ |
| | Time to Max<br>Depolarization Rate | **0.001** | $F(1,29) = 13.80$,<br>$\eta_p^2 = 0.322$ | 0.749 | $F(1,28) = 0.11$,<br>$\eta_p^2 = 0.004$ |
| | Time to peak | 0.961 | $F(1,29) < 0.01$,<br>$\eta_p^2 = 0.000$ | 0.265 | $F(1,28) = 1.30$,<br>$\eta_p^2 = 0.044$ |
| | Peak Amplitude | **0.028** | $F(1,29) = 5.34$,<br>$\eta_p^2 = 0.156$ | 0.110 | $F(1,28) = 2.73$,<br>$\eta_p^2 = 0.089$ |
| | Maximum Repolarization<br>Rate | **<0.001** | $F(1,29) = 16.45$,<br>$\eta_p^2 = 0.362$ | 0.970 | $F(1,28) < 0.01$,<br>$\eta_p^2 < 0.001$ |
| | Average Repolarization<br>Rate | **<0.001** | $F(1,29) = 15.42$,<br>$\eta_p^2 = 0.347$ | 0.748 | $F(1,28) = 0.11$,<br>$\eta_p^2 = 0.004$ |
| | Time to max<br>Repolarization Rate | **<0.001** | $F(1,29) = 17.35$,<br>$\eta_p^2 = 0.374$ | 0.438 | $F(1,28) = 0.62$,<br>$\eta_p^2 = 0.022$ |
| | AP2 Halfwidth | **<0.001** | $F(1,29) = 18.04$,<br>$\eta_p^2 = 0.384$ | 0.915 | $F(1,28) = 0.01$,<br>$\eta_p^2 < 0.001$ |
| | AP2 90% Width | 0.229 | $F(1,29) = 1.51$,<br>$\eta_p^2 = 0.049$ | 0.868 | $F(1,28) = 0.03$,<br>$\eta_p^2 = 0.001$ |
| | Afterhyperpolarization | **0.001** | $F(1,29) = 13.42$,<br>$\eta_p^2 = 0.316$ | 0.059 | $F(1,28) = 1.91$,<br>$\eta_p^2 = 0.122$ |
| Average of AP3+ | Maximum Depolarization Rate | **0.001** | F(1,29) = 14.11, $\eta_p^2 = 0.327$ | 0.371 | F(1,28) = 0.83, $\eta_p^2 = 0.029$ |
| | Average Depolarization Rate | **0.001** | F(1,29) = 15.08, $\eta_p^2 = 0.342$ | 0.445 | F(1,28) = 0.60, $\eta_p^2 = 0.021$ |
| | Time to Max Depolarization Rate | **<0.001** | F(1,29) = 27.15, $\eta_p^2 = 0.484$ | 0.716 | F(1,28) = 0.14, $\eta_p^2 = 0.005$ |
| | Time to peak | 0.332 | F(1,29) = 16.21, $\eta_p^2 = 0.033$ | 0.440 | F(1,28) = 0.61, $\eta_p^2 = 0.021$ |
| | Peak Amplitude | **0.023** | F(1,29) = 5.76, $\eta_p^2 = 0.166$ | 0.098 | F(1,28) = 2.93, $\eta_p^2 = 0.095$ |
| | Average Repolarization Rate | **0.001** | F(1,29) = 14.89, $\eta_p^2 = 0.339$ | 0.607 | F(1,28) = 0.27, $\eta_p^2 = 0.010$ |
| | Time to max Repolarization Rate | **0.002** | F(1,29) = 12.02, $\eta_p^2 = 0.293$ | 0.453 | F(1,28) = 0.58, $\eta_p^2 = 0.020$ |
| | AP Multi 90% Width | **0.001** | F(1,29) = 12.34, $\eta_p^2 = 0.298$ | 0.759 | F(1,28) = 0.10, $\eta_p^2 = 0.003$ |
| Interspike Properties | ISI Slope | **0.016** | F(1,26) = 6.71, $\eta_p^2 = 0.205$ | 0.184 | F(1,25) = 1.86, $\eta_p^2 = 0.069$ |
| | ISI Standard Deviation | **0.029** | F(1,26) = 5.33, $\eta_p^2 = 0.170$ | 0.934 | F(1,25) = 0.01, $\eta_p^2 < 0.001$ |
| | ISI min | **0.001** | F(1,26) = 15.36, $\eta_p^2 = 0.371$ | **0.001** | F(1,25) = 10.97, $\eta_p^2 = 0.305$ |
| | ISI max | **0.014** | F(1,26) = 6.93, $\eta_p^2 = 0.210$ | 0.625 | F(1,25) = 0.25, $\eta_p^2 = 0.010$ |
| | ISI covariance | 0.114 | F(1,26) = 2.67, $\eta_p^2 = 0.093$ | 0.417 | F(1,25) = 0.68, $\eta_p^2 = 0.027$ |
| | Instantaneous Frequency Median | **<0.001** | F(1,26) = 18.77, $\eta_p^2 = 0.419$ | **0.002** | F(1,25) = 11.41, $\eta_p^2 = 0.313$ |
| | Instantaneous Frequency Mean | **0.002** | F(1,26) = 12.30, $\eta_p^2 = 0.321$ | **0.003** | F(1,25) = 9.07, $\eta_p^2 = 0.266$ |
| | Instantaneous Frequency Max | **<0.001** | F(1,26) = 18.95, $\eta_p^2 = 0.422$ | **<0.001** | F(1,25) = 21.82, $\eta_p^2 = 0.466$ |
| | Instantaneous Frequency Min | **0.002** | F(1,26) = 11.28, $\eta_p^2 = 0.304$ | 0.335 | F(1,25) = 0.97, $\eta_p^2 = 0.037$ |
| Ratios | AP2/AP1 Halfwidth | 0.237 | F(1,26) = 1.47, $\eta_p^2 = 0.053$ | 0.492 | F(1,25) = 0.49, $\eta_p^2 = 0.019$ |
| | AP3+/AP1 Halfwidth | 0.342 | F(1,26) = 0.94, $\eta_p^2 = 0.035$ | 0.668 | F(1,25) = 0.19, $\eta_p^2 = 0.007$ |
| | AP2/AP1 AHP Amplitude | 0.463 | F(1,26) = 0.58, $\eta_p^2 = 0.022$ | 0.916 | F(1,25) = 0.01, $\eta_p^2 < 0.001$ |
| | APmulti/AP1 AHP Amplitude | 0.669 | F(1,26) = 0.19, $\eta_p^2 = 0.007$ | 0.387 | F(1,25) = 0.19, $\eta_p^2 = 0.030$ |
| | ISI max/min | 0.453 | F(1,26) = 0.58, $\eta_p^2 = 0.022$ | 0.113 | F(1,25) = 2.70, $\eta_p^2 = 0.097$ |

**Table 10.** Manual measurements positively correlate with script-derived values. We validated the script-derived values by randomly selecting 3 recordings for validation by using all 3 of their timeslots. Therefore an N = 9 files were analyzed in the script and with manual measurements using Clampfit 10.7. In all 7 measurements, values derived with the script were significantly and positively correlated with manual measurements. Values compared with a bivariate correlation and tested for significance (two-tailed; α = 0.05). Data analyzed in this table can be found in Supplementary File 1 “Validation” workbook.

| Measure | Correlation |
| --- | --- |
| **RMP** | $r(7) = 0.998, p < 0.001$ |
| **AP Threshold** | $r(7) = 0.982, p < 0.001$ |
| **Number of APs** | $r(7) = 1.000, p < 0.001$ |
| **ISI** | $r(7) = 0.999, p < 0.001$ |
| **Halfwidth** | $r(7) = 0.963, p < 0.001$ |
| **Maximum AP Repolarization Rate** | $r(7) = 0.998, p < 0.001$ |
| **AHP Amplitude** | $r(7) = 0.921, p < 0.001$ |

Our data demonstrates that the μOR modulates AP parameters, including the halfwidth and maximum rates of repolarization to suppress the excitability of interneurons. We observed halfwidth changes of 12% (S-responder mean) 21% (H-responder mean) and mean maximum repolarization rates of 25% or 40% (Table 7) - with also a high degree of variation between neurons (Figure 9). The consequences of AP kinetics have been explored in models with large synapses by using real and pseudo APs to shown that kinetics have consequences for voltage-gated calcium channel (VGCC) activity, calcium entry, and neurotransmitter release (Augustine, 1990; Klein & Kandel, 1980; Llinas, Steinberg, & Walton, 1981). Computer simulations and developmental data from synapses suggest that most VGCCs are activated by an action potential, but further broadening prolongs the kinetics of the VGCCs, and therefore contributes to longer duration of Ca^2+^ entry (Borst & Sakmann, 1998; Geiger & Jonas, 2000; Sabatini & Regehr, 1997), Additional data from the calyx of Held have added that depolarization phases affect the number of VGCCs recruited, while repolarization affects their kinetics to influence the amount or duration of Ca^2+^ entry. Paired recordings have shown that both features can modulate the release of neurotransmitter and the amplitude of postsynaptic potentials, depending on stages of development and adaptations at the synapse (Chao & Yang, 2019; Yang & Wang, 2006)

The αDTX-sensitive channels Kv1.1, Kv1.2, and Kv1.6 have previously been linked to regulation of cortical interneurons. In the neocortex, they have been found to strongly influence firing through their localization at axon initial segments of interneurons and excitatory neurons (Bekkers & Delaney, 2001; Guan et al., 2006). The have previously been linked to the excitability of cortical interneurons by modulating AP threshold and near-threshold spike frequency (Goldberg et al., 2008). In VIPergic interneurons, these currents affect the frequency of output (Porter et al., 1998). Genetic knockout of Kcna1 (encodes Kv1.1) and Kcna2 (encodes Kv1.2) are linked to seizures and ataxia in both humans and rat models (Adelman, Bond, Pessia, & Mayliet, 1995; Brew et al., 2007; Heeroma et al., 2009; Robbins & Tempel, 2012). Restoring regulated activity of these channels has been proposed as a mechanism underlying a treatment for Fragile X syndrome (Yang et al., 2020). Here, our data suggest that the μOR may upregulate these currents, as evidenced by αDTX’s attenuation of the DAMGO effect on firing frequency (Figure 9). The endogenous opioids of the neocortex are deeply involved in reward-seeking and motivated behaviors. Studies in animal models have shown enhanced μOR activity correlates with, and causes, binge-eating and drug intake (Blasio et al., 2014; Morganstern, Liang, Ye, Karatayev, & Leibowitz, 2012; Unterwald, Rubenfeld, & Kreek, 1994). The μOR is believed to reshape neocortical activity by releasing Pyramidal Neurons from inhibition by cortical interneurons (Férézou et al., 2007; Homayoun & Moghaddam, 2007; Jiang et al., 2019; H. K. Lee et al., 1980; Madison & Nicoll, 1988; Pang & Rose, 1989; Taki et al., 2000; Zieglgansberger et al., 1979). Yet despite the supporting evidence for μOR’s disinhibitory role in the neocortex, this receptor’s inhibitory influence has rarely been explored beyond its hyperpolarizing effects (Baldo, 2016; Férézou et al., 2007). Here, we found that the neocortical μOR has far-reaching inhibitory effects in neocortical interneurons that had not yet been described, which includes the potential upregulation of αDTX-sensitive channels to modulate interneuronal output through firing frequency.

## Materials and Methods

### Cell Cultures

All procedures were authorized by the Institutional Animal Care and Use Committee (IACUC) at University of Vermont. We dissected CD^®^ IGS Sprague-Dawley (Charles River) pregnant rat dams to harvest neocortical neurons from the frontal cortices of E21 rat embryos. Brain cortices were rinsed in Hibernate A (Gibco, Grand Island, NY), dissociated with papain (Worthington, Columbus, OH) and mechanically separated through gentle trituration with a pipette. Dissociated cortical neurons were and cultured on round 12mm glass, PEI-coated (Sigma-Aldrich, St. Louis, MO) coverslips at an approximate density of 3×10^4^ cells/cm^2^. We maintained the neurons in Neurobasal A, B27, Penicillin-Streptomycin and Glutamax (All from Gibco), in a humidified 37^°^ C incubator with 5% CO_2_. Cultures were transformed on day in vitro 1 with an AAV-mDlx-NLS-mRuby2 to induce expression of a red fluorescent protein in cortical interneurons. Culture media was half-replaced every 3 days. Experiments commenced between day in vitro 16-37. AAV-mDlx-NLS-mRuby2 was a gift from Viviana Gradinaru (Addgene viral prep # 99130-AAV1); http://n2t.net/addgene:99130;RRID:Addgene_99130).

### Whole-Cell Recordings

Cultured neurons were transferred from their growth media into a chamber perfused with ACSF (in mM: NaCl, 126; KCl, 5; NaH2PO4, 1.25; CaCl2, 2; MgCl2, 1; NaHCO3, 26; Glucose, 20; and pyruvate,5 (Férézou et al., 2007). The ACSF was warmed to 30°C and constantly bubbled with a mixture of 95% Oxygen and 5% CO2. When indicated, both CNQX (20 μM; Sigma-Aldrich) and dAP5 (50 μM; Sigma-Aldrich) were included in the ACSF, which was then constantly perfused through the recording chamber. Saline controls were done alternately with drug condition recordings throughout the course of the experiments. α-Dendrotoxin (αDTX) was purchased from Alomone Labs (Jerusalem, Israel).

Patch pipettes with resistances of 5-10MΩ were fabricated from borosilicate glass capillaries and filled with intracellular saline containing (in mM) K-gluconate, 144; MgCl2, 3; ethyleneglycol-bis(2-aminoethylether)-N,N,N′,N′-tetraacetic acid, 0.5; 4-(2-hydroxyethyl)-1-piperazineethanesulfonic acid, 10 (Férézou et al., 2007). The pH was adjusted to 7.2 with potassium hydroxide and confirmed for an osmolarity of 285/295 mosm. Whole-cell recordings were made with an Axopatch 200B (Axon Instruments) amplifier and Clampex 9.2.1.9 (Molecular Devices) software. Signals were sampled at 10kHz with a Digidata 1322a (Axon Instruments) DA converter. Passive membrane and AP kinetics measurements were automated using a custom Python program (cc_analysis.py) the source code for which is available at https://github.com/moriellilab/cc_analysis_46.

All 7 measures that we investigated in the hypotheses were manually tested-against script generated values. In all 7 cases, they significantly and positively correlated to values determined manually with the threshold peak-detector (see Supplemental Table S15) of Clampfit 10.7 (Molecular Devices). All analyses used measurements generated by the cc_analysis program, with the exception of spiking pattern, which was always determined manually.

### Current-clamp protocol

We selected neurons for recording that were fluorescent (red), reflecting their exposure to AAV-mDlx-NLS-mRuby2 during culturing. We adjusted membrane potentials for a junction potential of −11mV. Only neurons with a healthy appearance were selected for recording. We generally used neurons that were more hyperpolarized than −45mV for recording, though primarily the inclusion criterion was primarily the ability to fire APs from their natural RMP after the application of a depolarizing current.

A diagram of the experiment is illustrated on Figure 2. Upon acquiring a whole-cell configuration, we recorded for 3 minutes to ensure that the RMP was stable and more hyperpolarized than −45mV, and that the neuron was capable of firing APs with a depolarizing current from its RMP. The 20 current steps (1s duration) we used to elicit APs were adjusted for each recording during this waiting period, because neurons had slightly different RMPs, thresholds, and resistances that required modulating the current magnitude. Once calibrated for the neuron, we kept that setting constant for all 3 drug conditions for all timeslots for that neuron. We subsequently made the pre-drug recording, and then we applied these drugs through bath perfusion for 80s to ensure the concentration and response was stable. We then made the “post-DAMGO” (slot 2) recording after these 80s of incubation, then applied the next drug buffer for 80s (a combination of DAMGO and αDTX), and then made the post-αDTX+DAMGO recording (slot 3). Neurons were passively recorded from between timeslots for V_m_ changes, but these data were not considered in the analyses. Coverslips were discarded and replaced after a single use.

We quantified spontaneous APs by counting APs that occurred outside of the 1s current pulse. This comprised a noncontiguous period totaling 60 seconds in each timeslot. Spontaneous APs were not analyzed beyond counting them because most DAMGO-responders stopped firing spontaneous APs after DAMGO.

### Measurements of AP and membrane properties

APs were evoked with a 1s current pulse of varying amplitude. These pulses occurred 3s apart. Resting membrane potential (RMP) was determined from the median V_m_ measured during a 330 ms period immediately before the 1s current pulse. when the 1s current pulse was not being applied. This was composed of a noncontiguous period surrounding the current pulse, totaling 6.6s.

AP Threshold was determined from steady-state V_m_ at the first current application that induces an action potential. Mean interspike intervals were likewise derived from measuring the mean value for the space between spikes. Number of evoked APs was measured by averaging the number of APs also during episodes 15-18. The remaining parameters were likewise determined from the average value of episodes 15-18.

AP kinetic values (halfwidth and maximum repolarization rates) were also derived from episodes 15-18 of the 20 current steps. We averaged the values for APs 3-5 on each of the 4 current steps, and then averaged across the current steps to derive the final value. The halfwidth was determined as the width (in ms) of each AP at half the height of the AP.

### Data Display

The cc_analysis.py analysis program produced a Microsoft Excel spreadsheet with all measurements. To render the spreadsheet compatible with SPSS, we sorted the rows of recordings by filename and added in a row of column labels. SPSS syntax is compatible with the master spreadsheet included here. Additional columns were added to the excel spreadsheet that divided the post-DAMGO (or post-saline) scores by their initial (pre-drug) condition for each particular measure, which were ultimately graphed on box-and-whisker plots throughout this text. Every change in the post-DAMGO condition was some factor of 1 (initial) to show the fold change. Thus, the changes from baseline were determined for the DAMGO recordings and saline controls. While we had either 11 (H-responders) and 10 (S-responders), we randomly selected a group of comparable size to compare changes in the drug condition with entropic changes in the saline controls to identify the drug effect. In the box-and-whisker plots where a subset of 10 or 11 saline controls were required, we used Excel’s built-in random number generator function was used to sort the saline recordings for comparison groups.

### Statistics

Sample sizes were initially chosen under the expectation that only 5-20% of neurons would show a hyperpolarizing effect with DAMGO. We estimated that around 55 DAMGO recordings should enable us to record from ~10 DAMGO responders. We also recorded from 21 saline controls to enable us to compare the changes in DAMGO-exposed neurons to negative controls; groups are not of matching sizes because we expected that the whole DAMGO group would not be compared to the saline controls since responders would only compose some of the DAMGO-exposed neurons

Since we were examining for DAMGO or αDTX effects in 7 variables at a time, we first tested the combined variables in mixed-model repeated measures MANOVA tests first to mitigate the high (30.1%) family-wise error rate to an α = 0.05. If the omnibus test failed to produce significant results, we did not test each of the 7 variables individually (main hypotheses). However, we made exceptions for data in the post-hoc analyses of Table 9, since posthoc tests come with some assumptions of false positives.

All mixed-model repeated measures MANOVAs were conducted through IBM SPSS 27.0.0.0. We defined the independent variable as the timeslot (within-subject variable), and the between-subjects factor as the drug group (either saline or the DAMGO sequence). For these tests, we compared the 11 DAMGO H-responders (or 10 S-Responders) against all 21 saline-controls. For DAMGO effects we tested timeslot 1 and 2 data. For αDTX effects, we analyzed timeslot 2 and 3 data. We only reported Slot*Group interaction effects; the effect of Slot (time) most likely reflected entropic decay of the neurons over the course of the recording, while the effect of Group likely reflected stochastic differences of the neurons between the 2 groups of neurons – neither of which were a topic of these analyses. Therefore, in our experimental design, only the Slot*Group interaction was relevant to whether DAMGO or αDTX effects were present. In instances where the Slot*Group interaction was significant, we followed up by individually testing each of the 7 measures.

Post-hoc analyses (Table 9) were executed similarly with mixed repeated measures MANOVAs by testing sets of related variables to provide organizational and analytical grouping. APs analyzed in these tests were also measured in episodes 15-18, but first, second, and average (of APs 3-5) were analyzed separately. For instance, the first APs in episodes 15-18 were averaged across the episodes and analyzed separately from the other (second and later) APs in that episode. We excluded the 7 variables previously discussed in the main hypotheses to limit the posthoc analyses to data that were not previously analyzed. Results of their individual omnibus tests are noted in the figure legend for thoroughness, but we provided posthoc analytical data for individual with-subject contrasts regardless of whether their omnibus test was significant.

Statistical testing portrayed on the box-and-whisker plots and summary tables were conducted through in GraphPad Prism version 8.4.3 for Windows, GraphPad Software, San Diego CA, USA www.graphpad.com. Statistical testing on fold changes proceeded by first testing for skewness and kurtosis using D’Agostino-Pearson omnibus K2 test in GraphPad Prism. When the statistic was not significantly different from normal (*p* > 0.05), we conducted an unpaired t-test (one-tailed) to compare the fold changes in the DAMGO group with the saline group. When the distribution of either group was nonnormal (*p* < 0.05) we instead did the comparison with a one-tailed Mann-Whitney U-test.

