## Supplementary File Legends for "μ Opioid Receptors Modulate Action Potential Kinetics and Firing Frequency in Neocortical Interneurons"

Supplementary File 1: Master spreadsheet

The master spreadsheet (.xlsx) contains all script-derived numbers and ratios before they were copy/pasted to the appropriate figure or table throughout this text. It contains numerous workbooks intended to be used for different SPSS Syntax files. This spreadsheet was originally produced by the cc_analysis.py script and originally entitled "master_for_spss_PULSE_.xlsx" This file was subsequently reformatted into the file attached here to facilitate statistical analyses and consolidate data.

Workbook Directory:

**Main –** Master directory from which most other workbooks were derived. Green rows are S-responders. Yellow Are H-responders. Orange are saline controls. White rows are nonresponders.

**H_DAMGO** – Hyperpolarizers and saline controls. We used this workbook for posthoc tests for DAMGO, and to test the main hypotheses (7 measures) after DAMGO in H-responders.

**S_DAMGO –** S-responders and saline controls. We used to test DAMGO in S-responders.

**H_DTX** – H-responder DTX effects. Contains slots 2 and 3.

**S_DTX** – S-responders DTX effects.

**Nons_DAMGO** – Contains slots 1 and 2 from nonresponders and saline controls (to test for DAMGO effects in nonresponders

**Nons_DTX** – Contains slots 2 and 3 from nonresponders and saline controls (to test for DTX effects in nonresponders.

**Depolarizers** – Contains only information from the DAMGO depolarizers and saline hyperpolarizers (Figure 13).

**Validation** – Contains validation information used to for analyses in Table 10. This is the only workbook that is not a subset of "Main." Side-by-side columns are values from Clampfit or script. Alternating colors yellow and white separate each of the measures.

Supplementary File 2: SPSS Syntax

This file contains all SPSS Syntax run for posthoc tests and the main hypotheses. It is separated into several pieces. The first segment contains the SPSS Syntax to run the statistical comparisons on the main hypotheses. We ran this Syntax on: H_DAMGO for DAMGO effects in H-responders; S_DAMGO for DAMGO effects in S-responders; H_DTX for DTX effects on H-responders; S_DTX for DTX effects in S-responders.

The remaining segments were only run for posthoc tests. We ran these remaining Syntax codes on H_DAMGO for posthoc DAMGO effects, and H_DTX for posthoc DTX effects.
